# Allocation of resources among multiple daughter cells

**DOI:** 10.1101/2025.05.02.651883

**Authors:** Alison C.E. Wirshing, Roberto Alonso-Matilla, Michelle Yan, Samra Khalid, Analeigha V. Colarusso, David Odde, Daniel J. Lew

## Abstract

Cell division commonly produces two daughter cells, but there are many exceptions where large cells produce multiple daughters. Multiple fission of some green algae and bacteria, cellularization during embryogenesis of plants and insects, and growth of Ichthyosporeans, Chytrids, and Apicomplexans all provide variations on this theme. In some yeast species, a large multi-nucleate mother cell grows multiple buds (daughters) simultaneously. Here we address how mothers partition growth equally among their buds in the multi-budding yeast *Aureobasidium pullulans*. Bud growth is directed by actin cable networks that appear to be optimized for even partitioning despite complex cell geometries. Even partitioning does not rely on compensatory mechanisms to adjust bud volumes, but rather stems directly from effective equalization of polarity sites. These results reveal how conserved cell polarity and cytoskeletal networks are adapted to build complex morphologies in fungi.

## INTRODUCTION

Most cells proliferate by doubling their contents and then dividing in two. However, some cells grow considerably more, and then divide to form many daughters. For example, there are green algae and bacteria that can form large cells that give rise to many smaller daughters through synchronous multiple fission (Pląskowska et al., 2023; Ivanov et al., 2019; Chimileski et al., 2024). During plant and insect development, some embryos generate a large multi-nucleate cell that then divides into many, sometimes hundreds, of individual cells during a synchronous cellularization process (McCartney and Dudin, 2023). Analogous cellularization strategies are also employed by single-celled Apicomplexans (parasitic protists) (Francia and Striepen, 2014), Ichthyosporeans (relatives of animals) (Dudin et al., 2019), and Chytrids (relatives of fungi) (Medina et al., 2025). Among fungi, multinucleate hyphae can bud off multiple conidia (asexual spores) simultaneously, and some budding yeasts can produce multiple daughter cells in a single cell cycle through a process of multi-budding (Restrepo and Jiménez, 1980; Mims et al., 1988; Lubbehusen et al., 2003; Mitchison-Field et al., 2019; Henß et al., 2022). Producing multiple daughter cells requires that parents allocate resources appropriately to all daughters. How this is accomplished in different contexts remains an open question.

Here, we set out to address how growth potential is partitioned across multiple daughters by the multi-budding fungus, *Aureobasidium pullulans. A. pullulans* is a ubiquitous polyextremotolerant fungus that has long been cultivated for its commercial applications (Singh et al., 2008). More recently, *A. pullulans* has gained attention for its interesting growth strategy (Mitchison-Field et al., 2019; Petrucco et al., 2024; Wirshing et al., 2024; Goshima, 2022). Most yeasts, including model organisms like *Saccharomyces cerevisiae*, *Komagataella pastoris*, and *Kluyveromyces lactis*, as well as pathogens like *Candida albicans*, *Cryptococcus neoformans*, and *Ustilago maydis*, produce a single bud per cell cycle. Instead, *A. pullulans* mother cells are highly variable in size and can produce different numbers of buds (anywhere from 1 to >20) in a single cell cycle (Mitchison-Field et al., 2019; Petrucco et al., 2024; Wirshing et al., 2024). We have a deep mechanistic understanding of how buds grow from work in *S. cerevisiae*. The process begins with local activation of the conserved Rho-family GTPase Cdc42 to generate a single polarity site on the mother cell cortex (Pringle et al., 1995; Chiou et al., 2017). Cdc42 recruits the formin Bni1 to build actin cables oriented toward the polarity site, and myosin V motors deliver various cargos, including post-Golgi secretory vesicles, from the mother cell to the growing bud (Evangelista et al., 2002, 1997; Pruyne et al., 1998; Johnston et al., 1991; Sagot et al., 2002; Pruyne et al., 2004). Vesicle fusion is facilitated by Cdc42, thereby inserting new membrane and building new cell wall as the bud grows (Adamo et al., 2001). When there is only one bud, all relevant cargo traffics in a single direction. However, if a mother cell builds 2, or 3, or 17 buds, it is unclear whether or how a mother could ensure that every bud gets an equal share of its growth materials. One potential mechanism would be to monitor each bud’s growth and adjust delivery of materials in cases where buds grow too much or too little. Alternatively, daughter cells may be able to manage with widely varying inheritances from the mother by adjusting their behavior later on.

Here we find that *A. pullulans* mother cells grow sister buds at similar rates to reach similar sizes. As with *S. cerevisiae*, bud growth is driven by delivery of post-Golgi secretory vesicles from mother cells to each bud along actin cables. By perturbing actin and polarity machineries, we find no evidence for the existence of compensatory mechanisms acting to equalize sister bud growth. Instead, equitable partitioning of growth potential appears to depend on stringent equalization of the polarity sites that initiate and drive budding (Crocker et al., 2024). These findings provide insight into how an elaborate multi-polar cell shape is constructed.

## RESULTS

### *A. pullulans* equitably divides growth potential across multiple daughter cells

*A. pullulans* mother cells can vary over 70-fold in volume and contain variable numbers of nuclei (Petrucco et al., 2025; Wirshing et al., 2024). Larger cells contain more nuclei and tend to make more buds, which emerge from different positions on the mother cell surface (Wirshing et al., 2024; Mitchison-Field et al., 2019). Sibling buds growing from the same mother cell appeared similar in size, suggesting that bud size is regulated (**Figure 1A**). To measure bud volumes, we introduced a fluorescent cytosolic marker (three tandem copies of mCherry, 3xmCherry) and used confocal microscopy to capture entire cell volumes (**Figure 1B**). Quantification of bud volume variability confirmed that sibling buds had similar volumes (mean CV = 0.20), especially when mothers produced only two buds (mean CV = 0.12) (**Figure 1C**). This raised the question of how mothers ensure that their buds grow to similar sizes.

**Figure 1:**
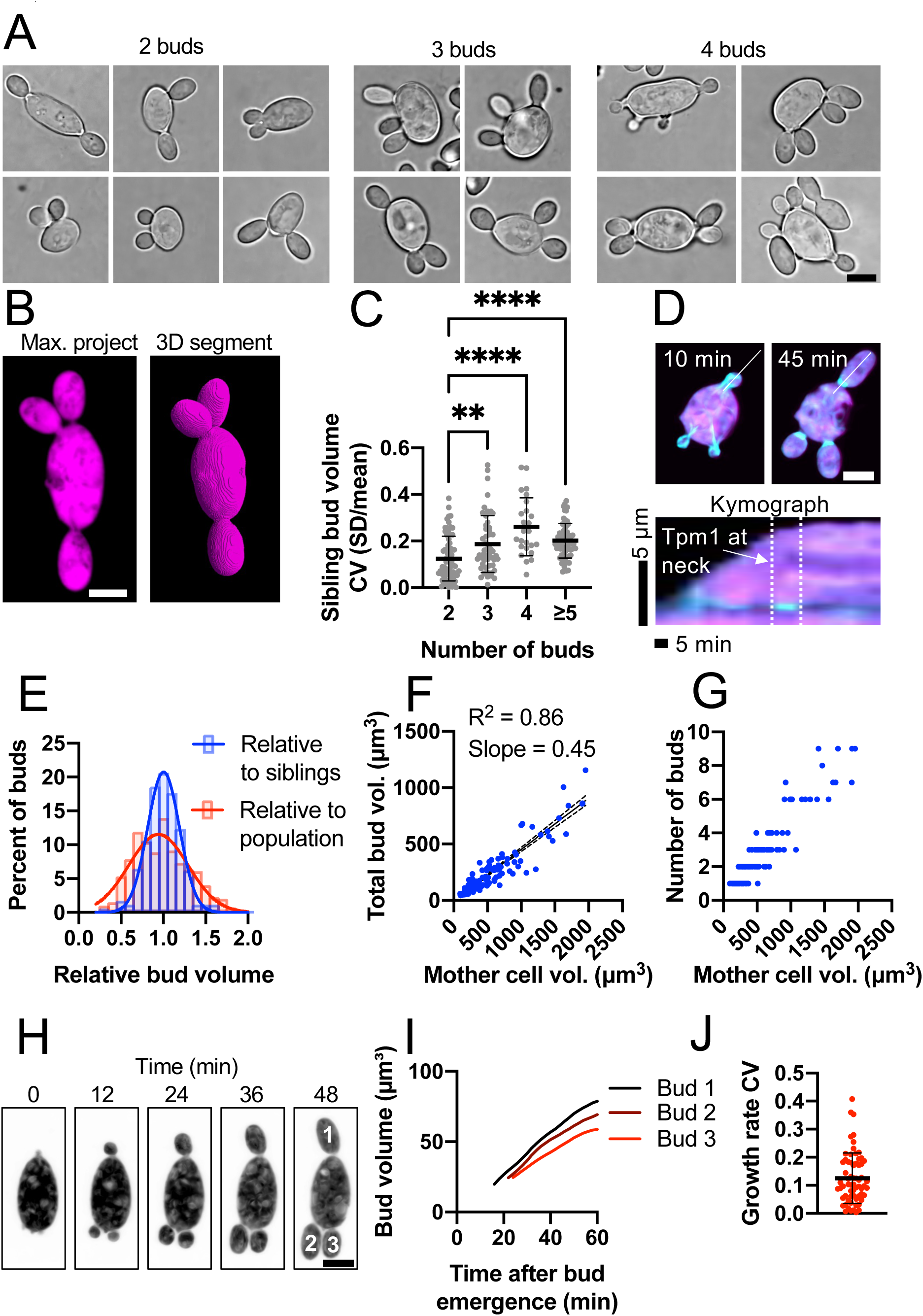
Bud size heterogeneity in *A. pullulans* multi-budding growth. (A) Brightfield images of mother cells (DLY24919) growing different numbers of buds from different positions. (B) Example confocal image of a cell expressing the cytosolic marker 3xmCherry and 3D segmentation used to measure cell volumes (DLY24919). (C) Coefficient of variation (CV) of bud volumes for mother cells making the indicated numbers of buds (n = 71, 53, 25, and 67 cells for 2-budded, 3-budded, 4-budded, and ≥5-budded categories). (D) Example maximum intensity projections of frames from a time series and a kymograph from a cell expressing mNG-Tpm1 and 3xmCherry (DLY25283). The white line indicates the line used to generate the kymograph. The dashed lines indicate when Tpm1 is at the neck. (E) Distribution of bud volumes relative to sibling buds (n = 339 buds) and relative to age-matched buds across the population with mNG-Tpm1 at the bud neck (n = 394 total buds). (F) Plot of mother cell volume and total volume of mature buds (with mNG-Tpm1 at bud necks) (n = 149 mother cells). (G) Plot of mother cell volume and the number of buds for the same cells shown in F. (H) Example maximum intensity projections of frames from a time series of a cell expressing 3xmCherry showing budding growth (DLY24919). (I) Time course of bud growth for the buds shown in H. (J) The CV of bud growth rates (n = 65 cells). Scale bars, 5 µm. The mean and standard deviation are shown in C and J. Statistical significance (C) calculated by one-way ANOVA (**, p ≤ 0.01; ****, p ≤ 0.0001).

A simple way to control bud size would be to have a set target size, such that buds below that size continue growing but once they reach the target size they stop growing. This would ensure that all buds in a population are borne at a set size. To assess final bud size at the time of cytokinesis, we expressed an N-terminal mNeonGreen (mNG)-tagged copy of *A. pullulans* tropomyosin at the *URA3* safe harbor locus (Wirshing et al., 2024). Expression of the probe did not compromise colony growth (**Supplemental Figure 1**). Tropomyosin decorates actin cables during bud growth and brightly decorates the cytokinetic ring at the bud neck only after buds have finished growing and are entering cytokinesis (**Figure 1D**). Using this probe to identify cells with mature (cytokinesis-stage) buds, we found that bud volumes varied much more across the population than they did when comparing siblings from the same mother (**Figure 1E**). Thus, there does not appear to be a universal bud target size. Instead, each mother equalizes sibling bud sizes.

We noticed that when similar-sized mother cells made different numbers of buds, those that made fewer buds grew larger buds than those that made more buds. This raised the possibility that the growth potential of a mother cell is set by its starting volume, and is divided by whatever number of sibling buds it produces. To test that idea, we plotted the sum of all sibling bud volumes against the mother volume (**Figure 1F**). Despite the large variation in mature bud sizes (**Figure 1E**) and bud numbers (**Figure 1G**), there was an excellent linear relationship between mother volume and the summed bud volume (**Figure 1F**), with a slope = 0.45. Thus, each mother grows daughters that sum to 45% of its starting size within a single cell cycle. Further, sibling buds emerged at similar times and grew at similar rates (**Figure 1H-1J**). Thus, sibling buds compete for a fixed mother growth potential that is equitably divided between them. To understand how this division of growth potential occurs, we next investigated the mechanisms driving *A. pullulans* budding growth.

### Secretory vesicles are produced throughout the mother cell volume and delivered to buds using F-actin

In fungi, polar growth is driven by local delivery and fusion of post-Golgi secretory vesicles carrying cell wall and membrane components (Pruyne et al., 2004). However, the organization of the secretory system differs between fungi. Some, like the yeast *K. pastoris*, have clustered endoplasmic reticulum exit sites (ERES) that give rise to Golgi stacks and local production of secretory vesicles (Papanikou and Glick, 2009). Others, like *S. cerevisiae*, lack Golgi stacks and produce secretory vesicles throughout the cell volume (Papanikou and Glick, 2009). The hyphal fungus *Aspergillus nidulans*, a closer relative of *A. pullulans*, also lacks Golgi stacks but ERES and Golgi cisternae are enriched at growing hyphal tips, suggesting that many vesicles are produced close to where they fuse (Pantazopoulou and Peñalva, 2009). If *A. pullulans* vesicles are produced locally within buds, then bud size equalization could reflect some mechanism to assign early secretory compartments fairly to each bud. On the other hand, if secretory vesicles are produced mainly in the mother, then bud size equalization would reflect some mechanism to direct even delivery of vesicles from the mother to each bud.

To visualize secretory compartments in *A. pullulans*, we fluorescently tagged conserved components of ERES (Sec13 and Sec23), early Golgi (Vrg4 and Anp1), late Golgi (Chs5 and Sec7), exocyst complex (Exo70, Sec3 and Sec8), or post-Golgi secretory vesicles (Sec4) (Methods) (Donovan and Bretscher, 2015; Zhu et al., 2019). With the exception of Sec4, all strains tags were on endogenous genes. These probes did not affect colony growth, except for Sec23-GFP which had a mild growth defect at 24°C (**Supplemental Figure 1**). ERES and Golgi compartments were distributed throughout the entire cell volume, as in *S. cerevisiae*. (**Figure 2A**). We conclude that post-Golgi secretory vesicles are produced throughout the mother cell volume, suggesting that they must be somehow delivered to buds. Consistent with such delivery, the exocyst and secretory vesicles accumulated at growing bud tips (**Figure 2B and 2C**). Rapid time-lapse imaging confirmed that GFP-Sec4 puncta (presumably vesicles) moved processively into growing buds (**Figure 2D).**

**Figure 2:**
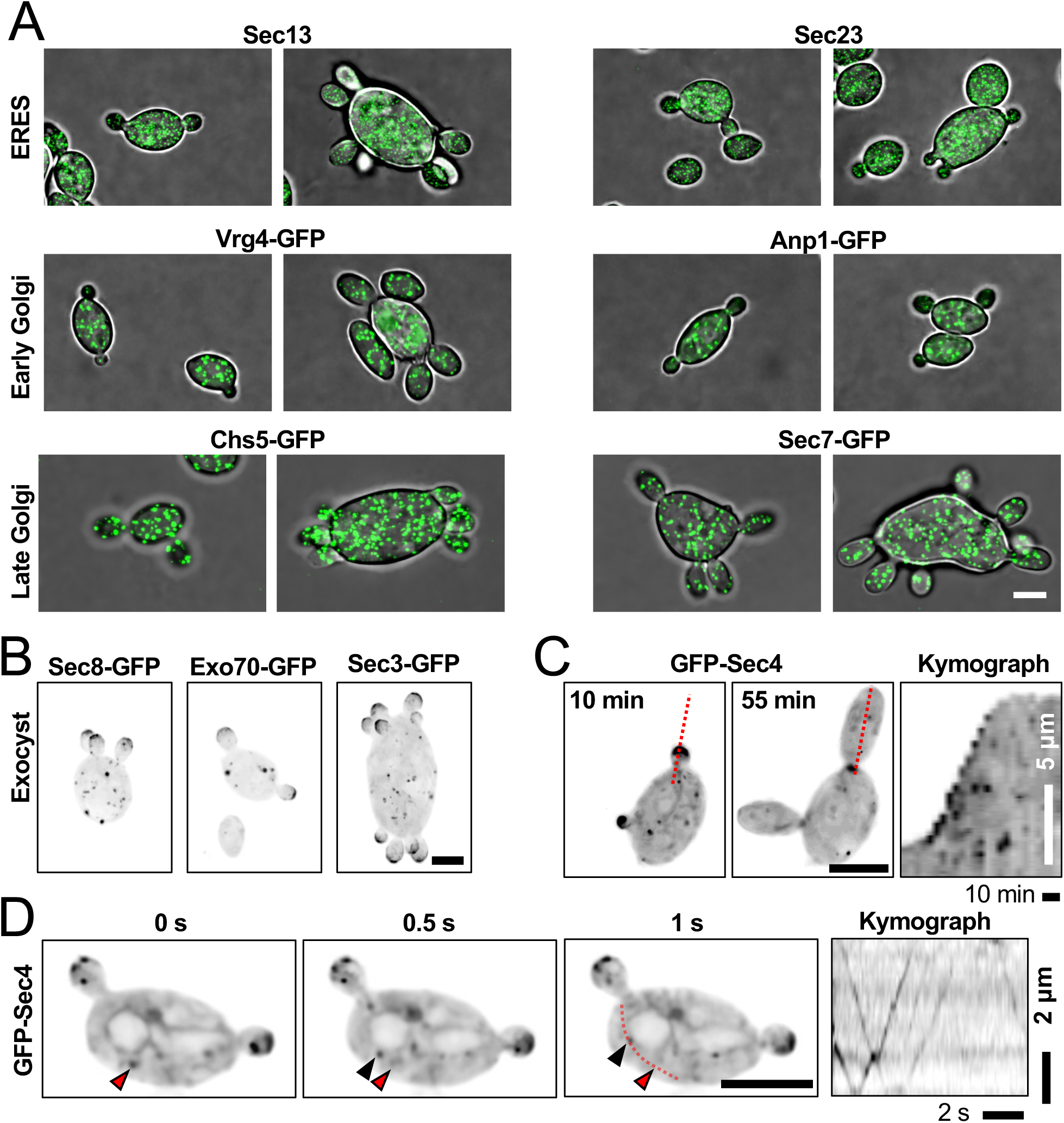
Organization of the *A. pullulans* secretory system. (A) Confocal maximum projection images of cells expressing the indicated markers for ER exit site (ERES) (DLY24715 and DLY24712), early Golgi (DLY24706 and DLY24709), and late Golgi (DLY24938 and DLY24935) (green). (B) Confocal maximum projection images of cells expressing the indicated markers for the exocyst complex (DLY25370, DLY25394, and DLY25367). (C) Maximum projection images and kymograph from time series of a cell expressing the secretory vesicle marker GFP-Sec4 (DLY24501). The dashed red line indicates the line used to make the kymograph. (D) Frames of a single-plane confocal movie highlighting processive vesicle movements. The red dashed line indicates the line used to make the kymograph. The top of the kymograph is toward the bud neck. Images in B-D are inverted to highlight dim signal. Scale bars, 5 µm.

To determine how secretory vesicles are delivered to buds, we tested the effects of depolymerizing F-actin or microtubules on bud growth. When cells were treated with benomyl to depolymerize microtubules, mitosis was disrupted but bud growth continued unimpeded (**Figure 3A**). Conversely, when cells were treated with Latrunculin B (LatB) to depolymerize actin, bud emergence and growth were blocked while the cells continued to grow isotopically and undergo mitosis (**Figure 3A**). We conclude that F-actin is required for polar growth in A. pullulans.

**Figure 3:**
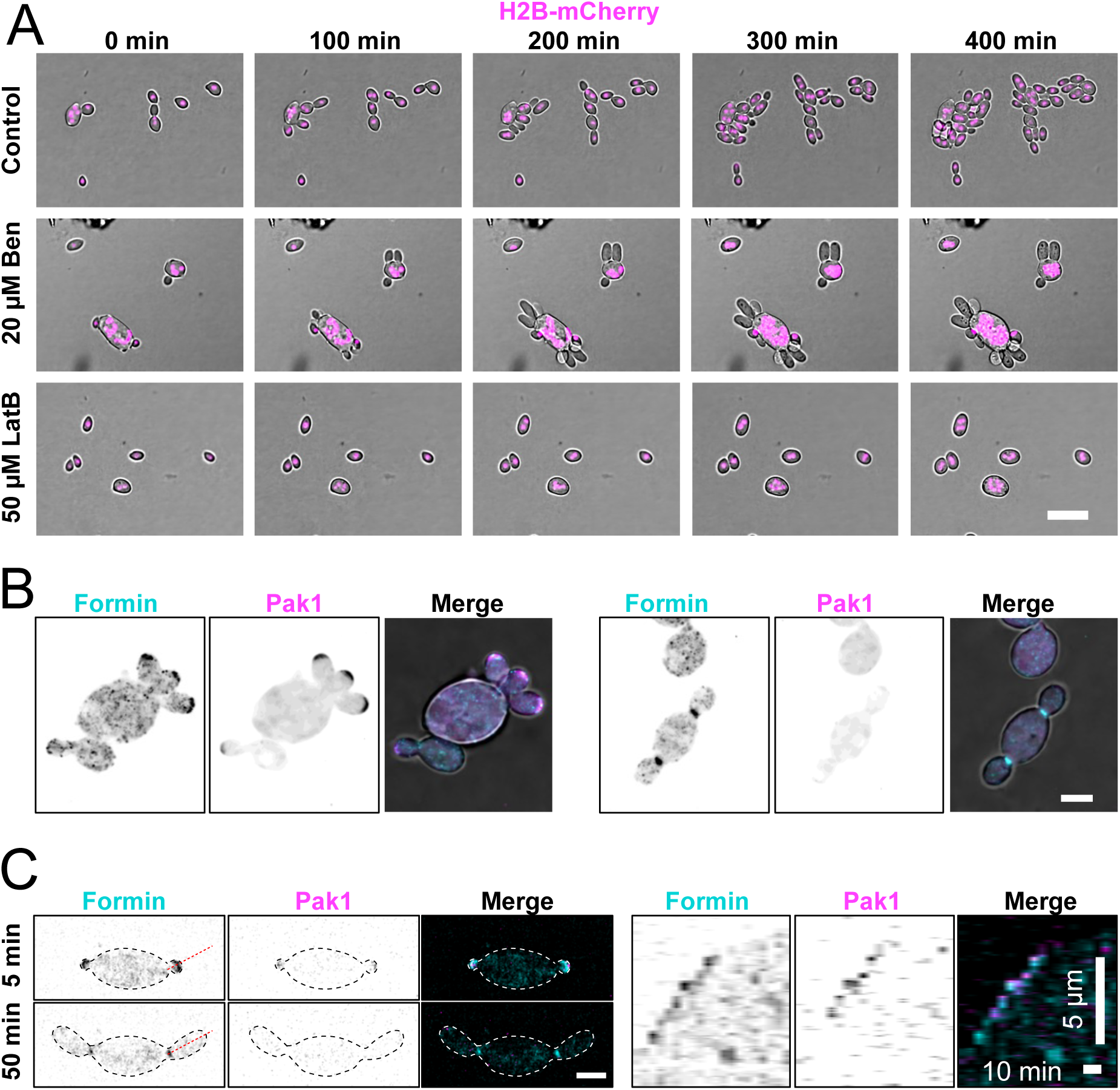
F-actin, and not microtubules, is required for bud growth. (A) Single-plane overlay bright-field and fluorescence images from time series of cells expressing the nuclear marker H2B-mCherry (magenta) (DLY24853) treated with LatB (50 µM), Benomyl (Ben, 10 µM), or vehicle control (DMSO). Scale bar, 10 µm. Start of movie (0 min) is ∼10 min after exposure to drug. (B) Representative confocal maximum projection images of cells expressing the polarity marker Pak1-mCherry (magenta) and fluorescent formin For1-3xmNG (cyan) (DLY25905). (C) Example maximum intensity projections of frames from a time series and associated kymograph from a cell expressing Pak1-mCherry and For1-3xmNG. The red line indicates the line used to generate the kymograph. Scale bar, 5 µm.

Fungal F-actin is commonly found in linear actin cables, associated with directed delivery of cargo, and cortical actin patches, associated with endocytosis (Moseley and Goode, 2006). Cables are assembled by a highly conserved family of proteins called formins (Sagot et al., 2002; Evangelista et al., 2002). We identified a single formin homologue in *A. pullulans* and C-terminally tagged it with three tandem copies of mNG (3xmNG). Tagging of the endogenous formin did not affect colony growth (**Supplemental Figure 1**). The *A. pullulans* formin colocalized with the polarity marker Pak1-mCherry (Crocker et al., 2024) at the tips of growing buds, and later in the cell cycle to the bud neck (**Figure 3B and 3C**). Thus, the formin appears positioned to nucleate a polarized network of actin cables in each bud. Together these results suggest that bud growth in *A. pullulans* is driven by the delivery of post-Golgi secretory vesicles from the mother cell to each bud along actin cables.

### Actin cable networks are dynamic and similar across sibling buds

Our finding that sibling buds grow at similar rates suggests that the post-Golgi vesicles supporting bud growth are delivered to each bud at similar rates. One way to achieve this is to ensure all sibling buds have comparable actin cable networks such that each network captures and delivers the same number of vesicles. To visualize actin cables, we stained cells with fluorescently tagged phalloidin. While phalloidin does not stain actin in wild-type *A. pullulans*, we previously showed that a single point mutation, *act1*^V75I^, is sufficient to confer phalloidin staining without altering cell growth or morphology (Wirshing et al., 2025). Phalloidin staining in the *act1*^V75I^ strain revealed actin cable networks extending from each bud into the mother, as well as actin patches clustered in each bud (**Figure 4A**). Surprisingly, the total length of each bud’s cable network was highly variable, frequently differing more than two-fold between sibling buds (**Figure 4A-4C**). If such uneven networks were stable during bud growth, we would expect some buds to harvest many more vesicles from the mother cell than their siblings, yielding large size differences. However, actin cables are often short-lived so the observed asymmetries might be transient. To test this, we monitored actin cables in live cells using the mNG-Tpm1 probe. Tropomyosin exclusively decorates formin-generated actin cable networks, allowing quantification of cables (Liu and Bretscher, 1989). To account for the cytosolic mNG-Tpm1 signal, we expressed cytosolic 3xmCherry in the same strain and subtracted the cytosol from the cable signal (See methods for details) (**Figure 4D**). Consistent with actin cables being dynamic, the mNG-Tpm1 signal in each bud varied over time (**Figure 4E**). While sibling buds occasionally had very different amounts of mNG-Tpm1, the asymmetry was transient and effectively averaged out over a 10-min window (**Figure 4F**). These results show that *A. pullulans* assembles a dynamic network of actin cables extending from each bud and that over time, sibling buds have similar networks.

**Figure 4:**
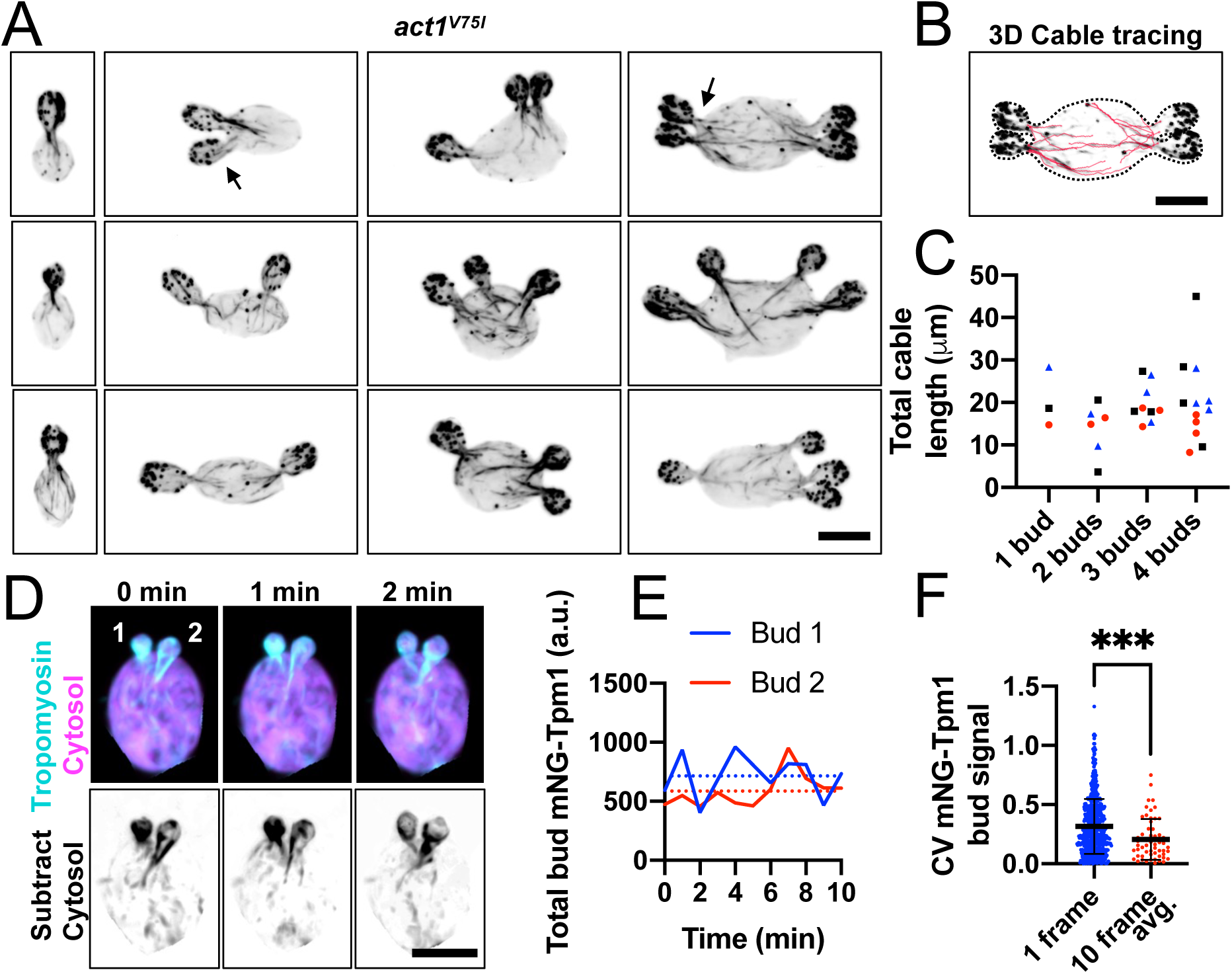
Actin cable networks in multi-budding cells. (A) Confocal inverted maximum projection images of *act1*^V75I^ cells fixed and stained with phalloidin (DLY24872). (B) Example of 3D cable tracing. Dashed line indicates cell outline. (C) Total cable lengths from each bud neck for the cells shown in E. For each category (number of buds), the same symbol is used for buds from the same mother. (D) Confocal merged maximum projection images from a time series of a cell expressing the cable marker mNG-Tpm1 (cyan) and cytosol marker 3XmCherry (magenta) (DLY25283). Inverted greyscale images show mNG-Tpm1 signal after subtracting the background cytosolic signal (see methods for details). (E) Plot of total mNG-Tpm1 signal in each bud over time from the cell in H. Dashed lines indicate the mean signal over 10 frames (10 min). (F) CV of mNG-Tpm1 signal among sibling buds for single snapshots (blue) or averaged across 10 frames (red) (n = 58 multi-budded cells). The mean and standard deviation are shown. Statistical significance calculated by two-way student’s T-test (****, p ≤ 0.0001).

### Does asymmetry in bud site position lead to a systematic difference in bud size?

If buds emerged at symmetric positions around the mother cell, then similar actin networks would suffice to generate similar-sized sibling buds. However, *A. pullulans* mother cells often produce sibling buds at quite asymmetric positions (**Figure 1A**). For example, if a mother cell produced two buds immediately adjacent to each other at one pole and another bud at the opposite pole, this inherent asymmetry in bud spacing might result in unequal growth. Perhaps competition for the local mother resources would yield fewer vesicles and slower growth for the adjacent buds than for the distant bud (**Figure 5A**). To gain an intuition for the degree of asymmetry that would be expected simply from asymmetric bud placement, we turned to a simple model for vesicle delivery (see Methods). The model assumes that: (i) Vesicles are produced at a constant rate throughout the mother volume; (ii) Vesicles diffuse in the cytoplasm until encountering and being captured by an actin cable; (iii) Actin cable networks emanate identically from each bud, with actin density decaying from the neck to the mother interior with a characteristic length scale; and (iv) The probability that a vesicle will hop on to a cable belonging to a particular bud’s network is proportional to the local actin density contributed by that bud.

**Figure 5:**
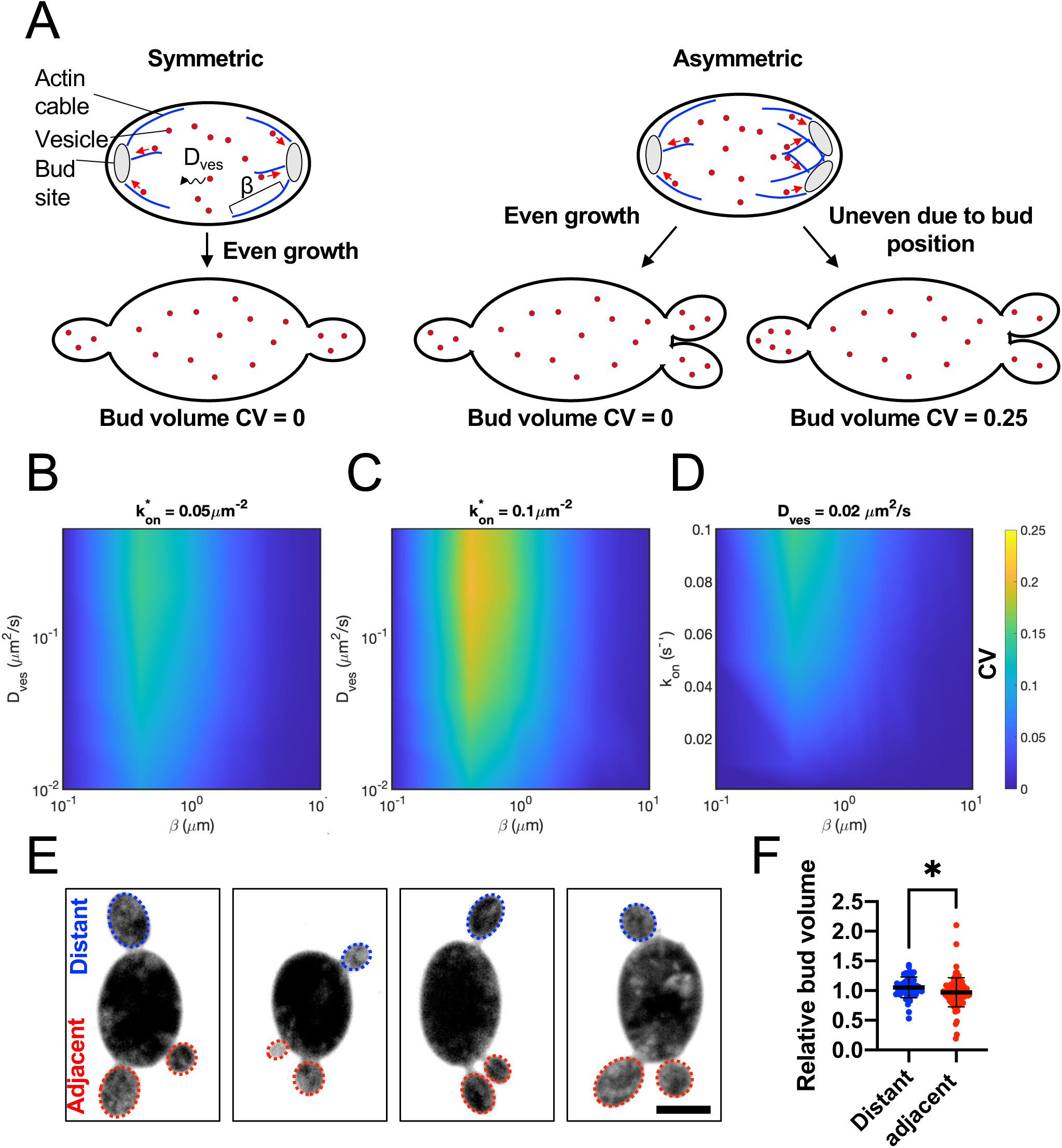
Effects of asymmetric bud placement. (A) Schematic showing key parameters of our model for vesicle behavior: actin cable number (affecting rate of vesicle capture k_on_), actin cable length scale (β) and vesicle diffusion constant (Dves). All buds are assumed to generate equivalent actin networks, so symmetrically placed buds capture vesicles at identical rates. However, asymmetrically placed adjacent buds may locally deplete vesicles where their actin networks overlap, leading to less vesicle capture than distant buds. (B-D) Calculated CV of vesicle delivery rate among asymmetrically placed sister buds as a function of model parameters. (E) Outliers: Maximum intensity projections of 3-budded cells expressing the cytosol marker 3xmCherry showing examples from the ∼10% of cells making the most asymmetric buds (DLY24919). (E) Bud volumes (relative to siblings) of three-budded cells comparing adjacent or distant buds (n = 49 distant and 98 adjacent buds). The mean and standard deviation are shown. Statistical significance calculated by two-way student’s T-test (*, p ≤ 0.05).

Using a mean field approximation, the model outputs a relative rate of vesicle delivery to each bud, which reflects differences in growth rates among the buds, and is assumed to correlate with their final sizes at maturity. There is no source of noise or stochasticity in the model, only the systematic difference that results from bud location. Symmetrically placed buds (**Figure 5A**, left) receive vesicles at identical rates, but asymmetrically placed buds (**Figure 5A**, right) differ, to a degree that depends on system parameters. Local vesicle depletion arises near each bud as vesicles are captured by actin cables, thereby slowing subsequent vesicle delivery. When buds are positioned close to one another, the overlap of their actin networks exacerbates depletion in the shared region. In this case, vesicle capture can outpace replenishment by diffusion, leading to a reduced local vesicle supply. This competition results in slower growth rates for adjacent buds compared to more distant ones. The magnitude of this effect is governed by the interplay of three key parameters: the vesicle diffusion constant, the characteristic length scale of the actin network, and the number of actin cables extending from each bud. We explored a range of values for these parameters informed by our observations and measurements from similar systems (see Materials and Methods). We find that with very few actin cables (low vesicle on-rate *k*on), any local depletion effects are small and bud asymmetry is very low (**Figure 5B**). With more actin cables, vesicle capture efficiency is increased, amplifying vesicle depletion especially in overlap regions between neighboring buds, thereby enhancing competitive asymmetries (**Figure 5C**). Depletion is mitigated when vesicle diffusion is very slow, because recruitment of vesicles by actin cables is diffusion limited. In this regime, buds receive comparable amounts regardless of spatial arrangement (**Figure 5C** bottom). With faster vesicle diffusion or high vesicle capture rates, the impact of actin network geometry becomes more pronounced (**Figure 5C,D**). For very short actin networks (i.e., a small characteristic length scale, with cables shorter than the inter-bud distance), vesicle depletion remains confined to the immediate vicinity of each bud neck, preventing competition between neighboring buds and resulting in equal vesicle recruitment rates across buds. At intermediate length scales (when actin cables from adjacent buds begin to overlap) the shared catchment region becomes more severely depleted than regions near isolated buds, leading to reduced vesicle recruitment for adjacent buds and increased size asymmetry (**Figure 5B-D**). This regime produces asymmetries comparable to those observed experimentally. At larger length scales, where actin cables extend more broadly into the mother cell, vesicle capture is distributed over a greater volume, reducing localized depletion and restoring more equal vesicle delivery across all buds (**Figure 5B-D**). Thus, depending on the specific vesicle diffusion, cable number, and actin network length scale parameters, the impact of asymmetric bud positioning on final bud size asymmetry can range from negligible to substantial.

Our measured bud volume variability (CV ∼0.2) could arise either from stochastic factors or from the systematic effects of asymmetric bud location. To address the latter, we focused on mother cells that grew two buds on one side (adjacent buds) and one bud on the other side (distant bud). While some of those cells had quite asymmetric bud volumes, bud size did not track systematically with bud location (**Figure 5E**). The distant bud was the largest only 41% of the time (20 out of 49 mother cells), not significantly different from the 33% that would be expected at random (p = 0.26, Chi-square test), and there was little difference in volume, on average, between distant and adjacent buds (**Figure 5F**). Thus, it appears that bud location is not a primary cause of bud volume variability.

### Mother cells do not appear to compensate for growth differences between buds

As we did detect rare instances in which apparently stochastic differences yielded significant bud volume variability, we wondered whether there were compensatory mechanisms to counteract initial differences and equalize bud sizes. For example, large buds might pause to allow smaller buds to catch up. Or, larger (smaller) buds might slow (speed) their growth rates to reach a final similar size (**Figure 6A**). To test for this, we monitored bud growth. When sibling buds were close in size, this was achieved by growing at similar rates without noticeable rate adjustments (**Figure 6B**). In outliers that produced particularly variable daughters, sibling buds grew at different rates throughout (**Figure 6C and 6D**). In some cells, multiple buds emerged at similar times but one bud initially grew slower than the others and later matched the growth rate of the faster siblings (**Figure 6E and 6F**). Across multiple mother cells, faster growing buds did not slow their growth and slower growing buds did not increase their growth to equalize bud volumes (**Figure 6G**). Thus, we do not detect obvious growth rate compensation.

**Figure 6:**
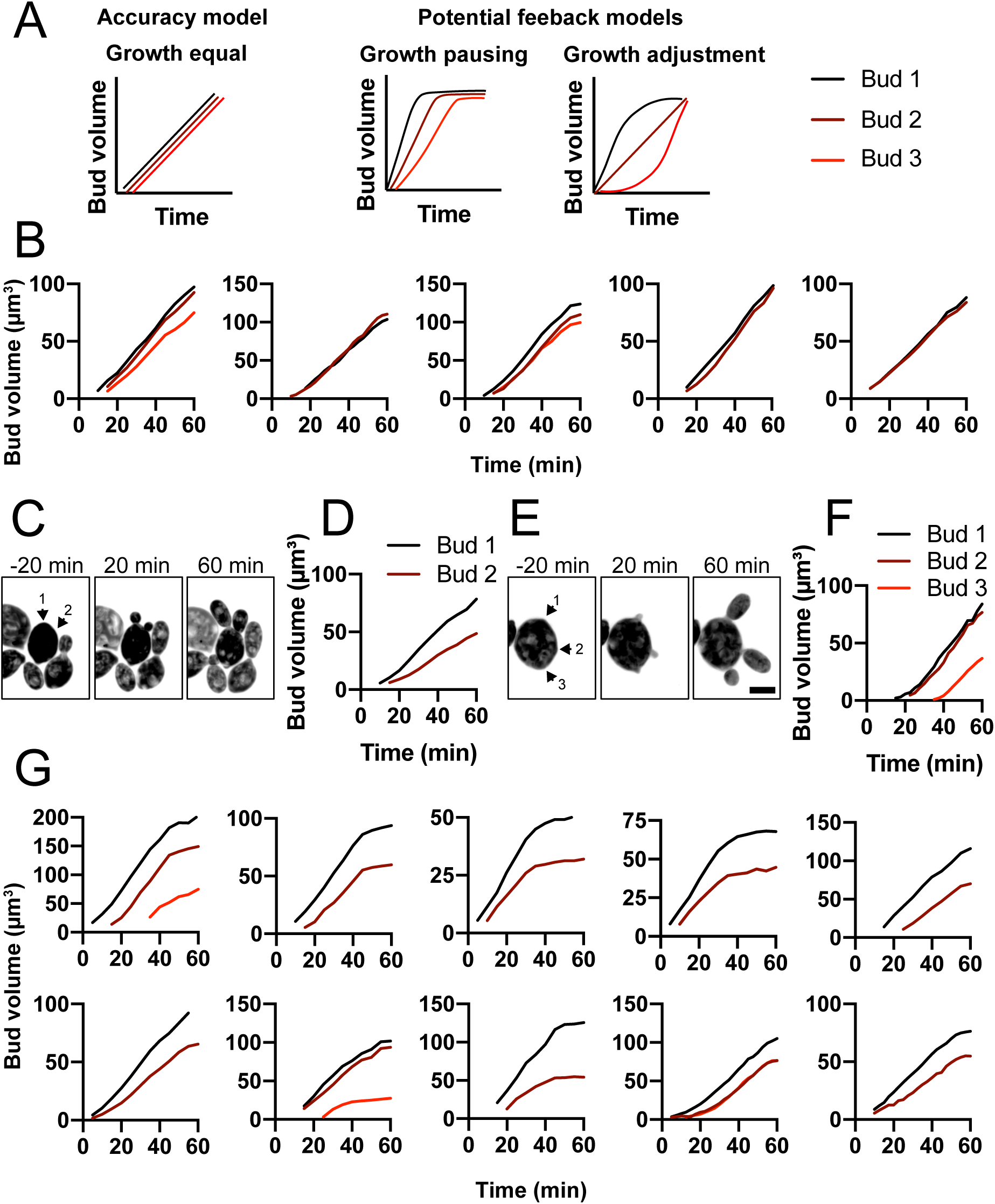
Mother cells do not adjust bud growth rates to equalize bud volumes. (A) Possible ways to grow buds similar in size. Either all buds grow similarly (accuracy model) or initial differences lead to compensatory changes (feedback models). (B) Plots showing segmented bud volumes from mother cells producing buds of similar sizes, representative of the large majority of cells. (C) Maximum projection images from time series of outliers making asymmetric buds (DLY24919). Arrows indicate bud emergence sites. (D) Bud volumes over time from cell in C. (E) Same as C showing a different example cell. (F) Bud volumes over time from cell in E. (G) Similar plots from other outliers.

As most mother cells produce daughters close in volume, uneven buds are rare. We reasoned that mildly perturbing vesicle delivery might increase bud variability, allowing us to detect compensation mechanisms that may simply have failed in the outliers. We found that cells treated with low doses (625 nM) of LatB continued budding even though they had few visible actin cables (**Supplemental Figure 2**). This treatment significantly increased bud volume variability, with many more mothers producing daughters dramatically different in volume (**Figure 7**). Even under these conditions, bud position did not predict bud volume, and distant buds were sometimes the smallest (**Figure 7A and 7C**). Thus, it is unlikely that the increased sibling bud volume variability with LatB treatment results from competition between adjacent siblings. Instead, other unknown differences between siblings may be exacerbated in this condition. Importantly, there was still no indication of growth rate compensation to equalize bud volumes (**Figure 7C-E**). The absence of compensation suggests that similarly-sized buds arise by accurately equalizing each bud’s actin network, rather than compensatory feedback between bud volume and growth rate.

**Figure 7:**
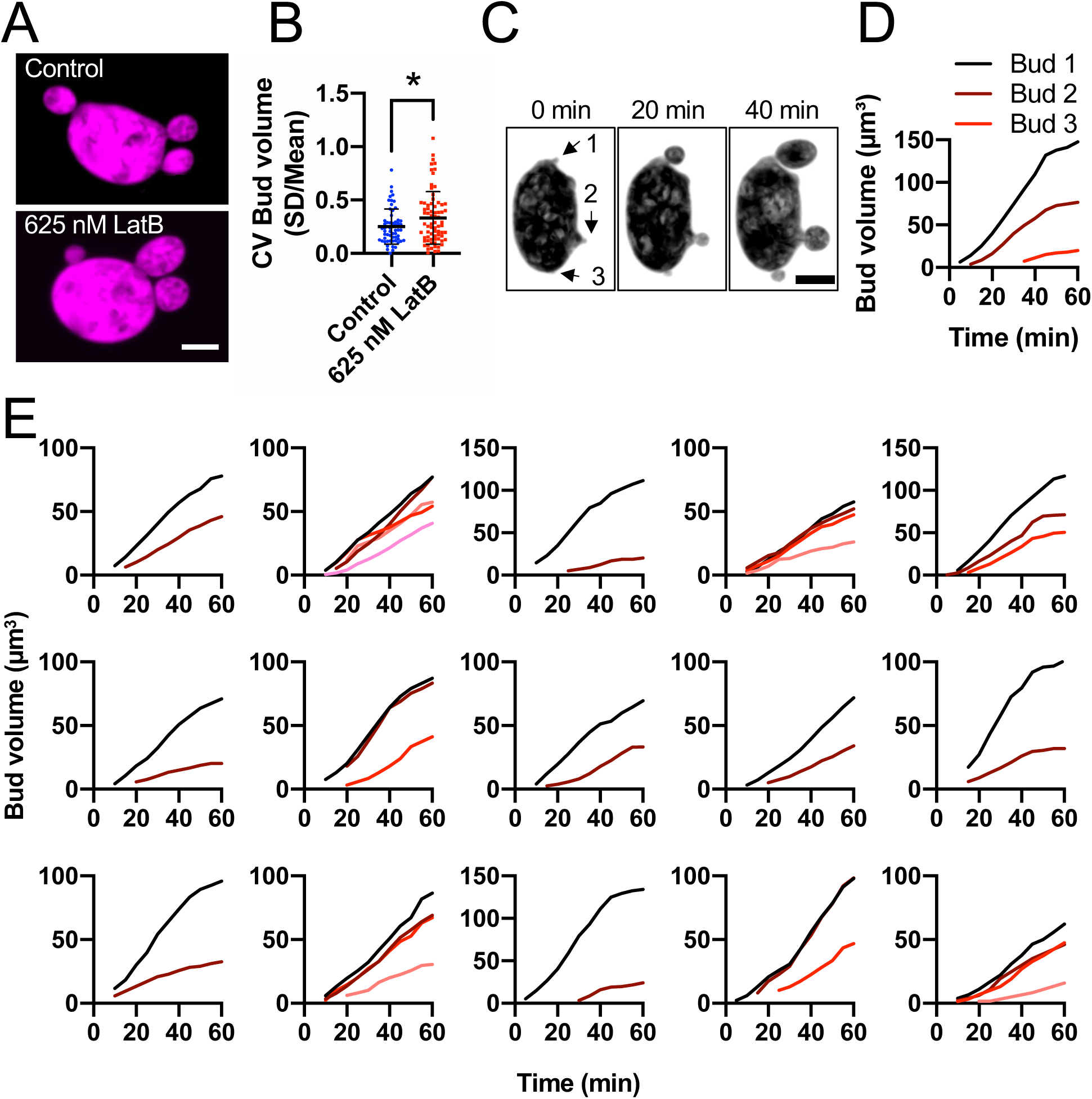
Perturbing F-actin increases bud volume variability. (A) Confocal maximum projection images of cells expressing the 3xmCherry cytosol marker (magenta) (DLY25647) treated with DMSO (control) or 625 nM LatB to perturb F-actin. Scale bar, 5 µm. (B) Bud volume CV is increased by LatB treatment (n = 58 control and 73 LatB treated cells). The mean and standard deviation are shown. Statistical significance calculated by two-way student’s T-test (*, p ≤ 0.05). (C) Confocal maximum projection images from time series of a cell expressing the cytosol marker 3xmCherry treated with 625 nM LatB. (D) Bud volumes over time from cell in C. (E) Similar plots from other examples of cells treated with 625 nM LatB.

### Equalizing polarity sites is required for even partitioning of growth potential

Budding in *A. pullulans* is initiated by the polarity regulators Cdc42 and Rac1 (Crocker et al., 2024). When cells make multiple buds, the polarity sites equalize their polarity factor contents, presumably serving as a necessary step to equalize each bud’s actin cable network. Polarity equalization requires the p21-activated kinase Pak1, which is recruited by active Rac1 (Crocker et al., 2024). By deleting the C-terminal kinase domain of Pak1 and replacing it with GFP, we were able to track localization of the N-terminal Rac1-binding domain (presumably reporting the location of active Rac1) in cells lacking Pak1 kinase activity. Cells expressing Pak1ΔC-GFP frequently produced buds of different sizes (**Figure 8A and 8B**), and sibling bud volumes were correlated with the amount of Pak1ΔC-GFP at bud tips (**Figure 8C**). Thus, the Pak1 kinase domain is needed to equalize polarity sites, and unequal polarity sites lead to growth of unequal buds. As with low-dose LatB treatment, we did not detect any evidence for compensatory mechanisms to equalize bud growth when initial growth rates differed (**Figure 8D-8F**). These results indicate that equalization of polarity factors at each polarity site is critical for even partitioning of growth potential.

**Figure 8:**
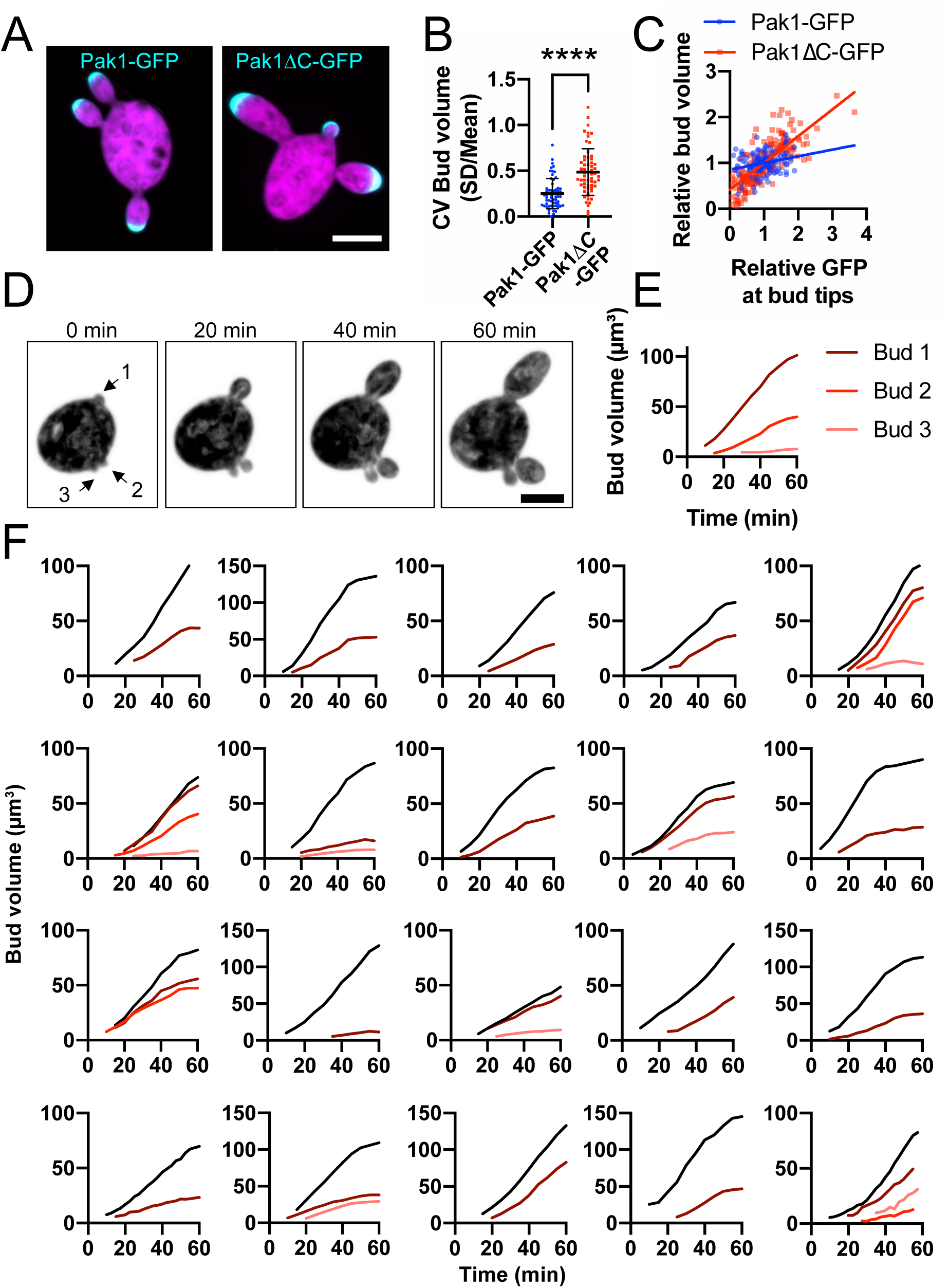
Equalizing polarity sites is required for even partitioning of growth potential. (A) Confocal maximum projection images of cells expressing the 3xmCherry cytosol marker (magenta) and Pak1-GFP (DLY25647) or Pak1 without the C-terminal kinase domain, Pak1ΔC-GFP (cyan) (DLY25650). Scale bar, 5 µm. (B) Bud volume CV is increased in cells expressing Pak1ΔC-GFP (n = 58 Pak1-GFP and 59 Pak1ΔC-GFP cells). The mean and standard deviation are shown. Statistical significance calculated by two-way student’s T-test (****, p ≤ 0.0001). (C) Plot of bud volume and total GFP signal (relative to siblings) at bud tips (Pak1-GFP or Pak1ΔC-GFP). Linear regression R^2^ = 0.07 for Pak1-GFP and 0.6 for Pak1ΔC-GFP. (D) Confocal maximum projection images from a time series of a *pak1*ΔC cell expressing the cytosol marker 3xmCherry (magenta) (DLY25650). (E) Bud volumes over time from cell in D. (F) Similar plots from other examples of *pak1*ΔC cells.

## DISCUSSION

The multi-budding yeast *A. pullulans* has an unconventional mode of proliferation in which mother cells of quite different sizes harbor varying numbers of nuclei and make variable numbers of buds. Here we characterized how growth potential was distributed across sibling buds in *A. pullulans*. We found that new-born buds varied in size considerably across a population of cells. However, mother cells of all sizes reproducibly constructed buds whose volumes added up to 45% of the mother cell volume. This suggests that rather than growing buds to a specific target size, mother cells produce a fixed proportion of their contents during a given budding cycle and divide those contents among the variable number of buds that are made. In most cases, the partitioning of this growth potential between sibling buds was quite even, with CVs of around 0.2. In principle, equal partitioning could arise in one of two ways. First, each bud might receive equal endowments through accurate partitioning from birth. Second, any stochastic or systematic differences between buds might be corrected by compensatory pathways. Our findings revealed an absence of compensation, supporting the hypothesis that the machinery for bud growth is accurately divided between all sibling buds from the onset of bud growth.

To probe the mechanisms of partitioning, we dissected how secretory compartments, cytoskeletal networks, and polarity machinery contribute to bud growth in *A. pullulans*. In *S. cerevisiae*, the post-Golgi secretory vesicles that drive bud growth are delivered from the mother cell to the bud along actin cables (Pruyne et al., 2004). Multi-budding growth in *A. pullulans* was also actin-dependent, and the organization of the *A. pullulans* secretory system and actin cable networks mirrored those of *S. cerevisiae* (Papanikou and Glick, 2009). Thus, the same machinery can direct growth to multiple buds simply by changing the number of polarity sites generated at the most upstream step.

Our central hypothesis is that similar polarity sites generate similar actin cable networks that draw secretory vesicles from the mother cell into each bud at similar rates. This is supported by our finding that increasing the variability of sibling bud polarity sites led to persistent asymmetric growth. However, two observations appeared to challenge that hypothesis. First, high-resolution snapshots of the actin cable networks in fixed cells revealed that sibling buds could display quite different cable networks. Second, bud emergence locations were often asymmetric, suggesting that even with similar actin networks, different buds would access distinct effective catchment areas in the mother. However, closer examination supported the equal polarity hypothesis.

Although instantaneous actin cable distributions could be quite asymmetric, lower-resolution live-cell imaging with a tropomyosin probe suggested that the cable networks are remodeled on a rapid timescale, leading to symmetric actin cable networks on a time-averaged basis. Asymmetric bud location provided a greater challenge, in that one might expect mothers to have difficulty delivering equal numbers of vesicles to buds in different geometries. However, simulations revealed that similar vesicle delivery across all buds could be achieved over a wide parameter space. The only exception was when buds made dense, locally overlapping actin cable networks causing vesicle depletion near adjacent buds. Under these conditions, buds growing close together on the mother cell surface would compete for vesicles, and grow smaller than distant sibling buds. However, bud position was a poor predictor of bud size in cells, suggesting that actin cable networks do not overlap to the correct degree to cause effective local vesicle depletion.

If our central hypothesis is correct, then the key to even partitioning of growth is the establishment of multiple equal polarity sites. A recent analysis of core polarity factors in *A. pullulans* suggested that polarity sites are indeed similar to each other on a time-averaged basis, and that this equalization requires the Rac1 effector, Pak1 (Crocker et al., 2024). Consistent with that view, we found that deletion of the kinase domain of Pak1 increased the variability in the amount of active Rac1 at each growing bud tip, leading to uneven bud growth.

How cells can equalize different polarity sites is an open question that has been addressed mainly through mathematical models of polarity circuits (Weyer et al., 2023; Goryachev and Leda, 2017; Chiou et al., 2018, 2021; Jacobs et al., 2019; Brauns et al., 2021). When multiple polarity sites are present, they can display one of three interactions. Competition, analogous to coarsening or Ostwald ripening phenomena in physics, yields a single “winning” polarity site. Competition for limiting cytoplasmic polarity factors is driven by the local positive feedback required to build a polarity site in models that assume mass conservation of polarity factors (Chiou et al., 2017; Goryachev and Leda, 2017). Coexistence occurs when the competition timescale is long relative to the relevant biological timescale (Chiou et al., 2018). Both competition and coexistence yield unequal polarity sites. Only the third scenario, equalization, yields equal polarity sites. This behavior arises in models that relax the mass conservation assumption (Weyer et al., 2023; Brauns et al., 2021), or add negative feedback or other features on top of the core positive feedback (Chiou et al., 2021; Howell et al., 2012; Jacobs et al., 2019). In a possibly related phenomenon, global positive feedback through a shared activator was recently proposed to promote equalization of centromere sizes in a shared cytoplasm (Banerjee and Banerjee, 2025). In *A. pullulans*, Pak1 has been suggested to promote polarity site equalization by providing negative feedback (Crocker et al., 2024), though a mechanistic understanding of what Pak1 does is currently lacking.

After initiating growth of multiple buds, *A. pullulans* mother cells presumably allocate organelles across all siblings. While some organelles (e.g. vacuole) may be generated *de novo* in daughters, others (e.g. mitochondria and nucleus) cannot and presumably rely on delivery from the mother cell (Weisman et al., 1990). *S. cerevisiae* delivers organelles to the bud using actin motors (Pruyne et al., 2004), and tightly regulates organelle inheritance to ensure that the bud receives, and the mother cell retains, the correct amount of each organelle (Knoblach and Rachubinski, 2015; Obara et al., 2024; Weisman, 2006; Westermann, 2014; Fagarasanu et al., 2010). How such mechanisms may be adapted to the more complicated geometry of a multi-budding cell is unclear. Are there mechanisms to ensure that siblings inherit similar organelle contents? Alternatively, do daughters compensate for differences by altering organelle biogenesis/autophagy rates? Organelle allocation in multi-polar cells remains a fascinating mystery.

Multi-polar growth is not unique to *A. pullulans*. Hyphal branching fungi, neurons, and plant xylem and epidermal cells all exhibit multi-polar growth and use this to build their unique shapes (Knechtle et al., 2003; Igisch et al., 2022; Oda and Fukuda, 2012; Dotti et al., 1988; Sugiyama et al., 2017). Additionally, there are other multi-budding fungi. The *Aureobasidium*-related *Paracoccidioides* species also undergo multi-budding growth, as do the very distantly related *Mucor* species (Lubbehusen et al., 2003; Restrepo and Jiménez, 1980). It may be that multi-budding growth enables a rapid dispersal of vegetative cells. Indeed, in *Paracoccidioides* and *Mucor* species multi-budding growth is thought to help spread fungal cells within mammalian hosts during infection (Aristizabal et al., 1998; Lee et al., 2013; Li et al., 2024). For *A. pullulans*, multi-budding growth may improve dispersal in aquatic environments (Mitchison-Field et al., 2019). Similar multi-polar structures are used throughout the fungal kingdom to facilitate spore formation and dispersal (Dijksterhuis, 2019). For example, ascomycetes such as *Aspergillus* form elaborate conidiophores that give rise to thousands of asexual spores called conidia that all grow from several branches (metula) off a single large cell (vesicle) (Mims et al., 1988). Basidiomycete fungi produce sexual spores, basidiospores, that bud four or eight at a time from a single large cell (basidium) (Campos and Costa, 2010; Kües, 2000). The fact that multi-polar structures are ubiquitous throughout the fungal kingdom highlights their utility. It also suggests that tuning of a highly conserved polarity machinery can produce different polarity patterns enabling cells to grow into diverse shapes. Thus, insights into polarity patterning and resource partitioning gleaned from *A. pullulans* will likely translate to a broader understanding of fungal morphogenesis.

## METHODS

### *A. pullulans* strains and maintenance

Strains used in this study are listed in **Supplemental Table 1**. All experiments were conducted with derivatives of *Aureobasidium pullulans* strain EXF-150 (Gostinčar et al., 2014). Unless otherwise indicated, *A. pullulans* was grown at 24°C in standard YPD medium (2% glucose, 2% peptone, 1% yeast extract) with 2% BD Bacto^TM^ agar (214050, VWR) in plates.

### Plasmid design and molecular biology

To express the cytosolic marker, 3xmCherry, a plasmid (DLB4820) was constructed with three tandem copies of codon-optimized mCherry under control of the Chytrid *S. punctatus* histone H2A promoter flanked by *A. pullulans URA3* sequences that enable replacement of *ura3Δ::Hyg^R^* with *URA3* (Wirshing et al. 2024). DLB4820 was derived from pAP-U2-1 (Addgene # 236472) by removing the H2B promoter-driven 3xGFP (Colarusso et al., 2025).

To express the actin cable marker, mNG-Tpm1 (Tropomyosin, protein ID 317632), a plasmid (DLB4897) was constructed expressing codon-optimized mNG-linker-Tpm1 (linker as in (Dhar et al., 2024)) using the native *TPM1* promoter and terminator (1 kb flanking sequences), cloned next to a Hygromycin resistance cassette. The mNG, Hygromycin resistance cassette, and plasmid backbone sequences were from pAPInt-mNG-Hyg (Addgene # 236483) (Colarusso et al., 2025).

To express the nuclear marker, histone H2B, a plasmid (DLB4847) was constructed with the Chytrid *S. punctatus* histone H2B promoter driving expression of H2B-mCherry. DLB4847 was derived from DLB4815 (Wirshing et al., 2024)by removing the NLS-GFP sequence.

To express the secretory vesicle marker, GFP-Sec4 (protein ID 271553), a plasmid (DLB4775) was constructed expressing codon-optimized GFP-*SEC4* coding under control of the *scACT1* promoter. DLB4775 inserts *SEC4* and removes the histone H2B promoter and 3xmCherry in pAP-U2-2 (Addgene # 236473) (Colarusso et al., 2025).

To visualize the formin, For1 (protein ID 369820) was C-terminally tagged with three tandem copies of mNG using pAPInt-3xmNG-Hyg (DLB4992), generated by introducing two additional copies of mNG into pAPInt-mNG-Hyg (Addgene # 236483) (Colarusso et al., 2025).

All PCR fragments were amplified with Phusion™ Hot Start Flex 2X Master Mix (M0536L, NEB) following the manufacturer’s instructions, and plasmids were built using NEBuilder HiFi DNA Assembly Master Mix (E2621L, New England Biolabs). All plasmids were confirmed by whole plasmid sequencing (Plasmidsaurus, Eugene, OR, USA). A list of plasmids used in this study are in **Supplemental Table 2**.

### Preparation of linear DNA for transformation into *A. pullulans*

Linear transforming DNA was prepared by PCR or restriction digest. Plasmids DLB4820, DLB4897, DLB4847, and DLB4775 described above were linearized by restriction digest and used directly in yeast transformations. Three-part-PCR (Colarusso et al., 2025) was used to introduce a C-terminal GFP tag onto the native *SEC13* (protein ID 378285), *SEC23* (protein ID 324106), *VRG4* (protein ID 355218), *ANP1* (protein ID 343193), *CHS5* (protein ID 352483), *SEC7* (protein ID 269067), *SEC8* (protein ID 378094), *EXO70* (protein ID 346362), *SEC3* (protein ID 402877), or *PAK1* (protein ID 374035). Three-part-PCR was also used to introduce the C-terminal 3xmNG tag on *FOR1* (protein ID 369820). The 3-part-PCR protocol involves amplifying separate 1 kb flanking sequences and the GFP construct with selection cassette for each gene, using primers that introduce 60-70 bp overlap between the three PCR products. Flanking sequences were amplified from EXF150 genomic DNA and the GFP-drug resistance cassette was amplified from vectors pAPInt-GFP-Nat (Addgene # 236476), pAPInt-GFP-Hyg (Addgene # 236477), or pAPInt-3xmNG-Hyg (DLB4992). Each PCR product was amplified using Phusion™ Hot Start Flex 2X Master Mix (M0536L, NEB) in four to eight reactions. These reactions were then pooled and concentrated by adding 0.1 volume of 3 M NaOAc and 2.5 volumes of 100% ethanol followed by incubation on ice for 10 min. The DNA-ethanol mixture was then added to a silica DNA binding column (T1017-2, New England Biolabs), rinsed with 400 µl wash buffer (80% EtOH 20 mM NaCl 2 mM Tris pH 8), allowed to dry for 1 min, and then eluted in ∼30-60 µl 10 mM Tris pH 8 at a final concentration of ∼1 µg/µl. After concentrating, three individual PCR products (both 1 kb flanking sequences and the central construct) were added to *A. pullulans* competent cells at an equimolar ratio in a final volume of 15 µl. All primers used in strain construction are listed in **Supplemental Table 3**.

### *A. pullulans* transformation

*A. pullulans* was transformed using PEG/LiAc/ssDNA as described previously (Wirshing et al., 2024). Briefly, cells were grown to a density of ∼10^7^ cells/ml in YPD (4% glucose), harvested by centrifugation, and rinsed with sterile water followed by competence buffer (10 mM Tris pH 8, 100 mM LiOAc, 1 mM EDTA, 1 M sorbitol). Cells were resuspended in competence buffer in a final concentration of 2×10^9^ cells/ml in a final volume of 50 µl (∼10^8^ cells in each tube). At this stage, competent cells were stored in the –80°C freezer or used directly by adding 2.5 µl 40% glucose, 10 µl carrier DNA (10 mg/ml single-stranded fish-sperm DNA, 11467140001, Roche), 15 µl of transforming DNA, and 600 µl of transformation buffer (10 mM Tris pH 8, 1 mM EDTA, 40% (w/v) PEG 3350). Cells were incubated in transformation buffer for 1 hour at 24°C while rotating. Following the 1-hour incubation, 30 µl 40% glucose was added, and the cells were heat shocked for 15 min at 37°C. Cells were centrifuged and the transformation buffer was removed. For selection of prototrophs on media lacking uracil, cells were resuspended in 250 µl 1 M sorbitol and spread onto two doURA, (6.71 g/L BDDifcoTM Yeast Nitrogen Base without Amino Acids, BD291940, FisherScientific, 0.77 g/L Complete Supplement Mixture minus uracil, 1004-100, Sunrise Science Products, 2% glucose, and 2% BD Bacto^TM^ agar, 214050, VWR) plates. For antibiotic selection, cells were first allowed to recover by resuspending in 1 ml of 1 M sorbitol and adding to a culture tube with 1 ml YPD (2% glucose). Cells were recovered for 4-24 hours at 24°C with agitation at 100 rpm before plating. Transformants were selected on YPD plates supplemented with 70.4 µg/ml Hygromycin B (400051-1MJ, Millipore), or 50 µg/ml Nourseothricin (N1200-1.0, Research Products International).

### Live-cell imaging

For imaging experiments, a single colony was used to inoculate 5 ml of YPD (2% glucose). Cultures were grown overnight at 24°C to a density of 1-5×10^6^ cells/ml. Cells were pelleted gently at 9391 rcf for 10 s and resuspended at a final density of ∼7×10^7^ cells/ml. Approximately 2×10^5^ cells were mounted on a 8-well glass-bottomed chamber (80827, Ibidi) and covered with a 200 µl 5% agarose (97062-250, VWR) pad made with CSM (6.71 g/L BD Difco^TM^ Yeast Nitrogen Base without Amino Acids, BD291940, FisherScientific, 0.79 g/L Complete Supplement Mixture, 1001-010, Sunrise Science Products, and 2% glucose). All imaging was conducted at room temperature (20-22°C).

To measure cell volumes, cells expressing the cytosolic marker 3xmCherry were imaged on a Nikon Ti2E inverted microscope with a CSU-W1 spinning-disk head (Yokogawa), CFI60 Plan Apochromat Lambda D 60x Oil Immersion Objective (NA 1.42; Nikon Instruments), and a Hamamatsu ORCA Quest qCMOS camera controlled by NIS-Elements software (Nikon Instruments). For still images, the entire cell volume was acquired using 75 Z-slices (at 0.2 μm step intervals). Exposure times of 50 ms at 50% laser power (excitation 561 nm) were used. To monitor bud volume over time, Z-stacks (17 slices, 0.7 µm interval) were acquired every 5 minutes for 3 hours using 50 ms exposures at 15% laser power (excitation 561 nm). In strains also expressing the polarity marker, Pak1-GFP or Pak1ΔC-GFP, the 488 nm laser at 50% or 15% laser power was used for still and time-lapse imaging experiments, respectively, with 50 ms exposure time.

To monitor tropomyosin dynamics, cells expressing mNG-Tpm1 and 3xmCherry were imaged with the same setup as above at 1-or 5-minute intervals for 1 or 3 hours. The entire cell volume was captured using 17 Z-slices (0.7 μm step intervals). Exposure times 50 ms and 15% laser power were used to image mNG-Tpm1 (488 nm) and 3xmCherry (561 nm).

To characterize the secretory pathway, still images of cells expressing GFP-tagged components of the secretory pathway (Sec13, Sec23, Vrg4, Anp1, Chs5, Sec7, Sec8, Exo70, and Sec3) were imaged as above. The entire cell volume was acquired using 75 Z-slices (at 0.2 μm step intervals). Exposure times of 50 ms at 50% laser power (excitation 488 nm) were used. To image GFP-Sec4, stacks of 17 Z-slices (0.7 μm step intervals) were acquired every 5 minutes for 3 hours using 50 ms exposure at 15% laser power (488 nm). To capture rapid GFP-Sec4 labeled vesicle movements, a single slice was captured every 100 ms for 10 seconds using 50 ms exposure time and 50% laser power (488 nm).

Formin localization was determined by imaging cells expressing For1-3xmNG and the polarity marker Pak1-mCherry on the same spinning disk setup as above. Still images were acquired using Z-stacks (75 slices, 0.2 µm interval) with 50 ms exposure time and 50% laser power using the 488 nm and 561 nm lasers to excite For1-3xmNG and the polarity marker Pak1-mCherry, respectively. Formin dynamics during the cell cycle were captured by acquiring a Z-stack (17 slices, 0.7 µm interval) every 5 minutes for 3 hours using 15% laser power and 50 ms exposure time for both the 488 and 561 nm lasers.

### Benomyl and LatB treatment

The effect of actin or microtubule depolymerization was determined by treating cells with LatB (625 nM or 50 µM final concentration) or benomyl (20 µM final concentration), respectively. Drug stocks were prepared in DMSO. LatB (10 mM stock) or benomyl (20 mM stock) were added to the CSM agarose pads used for imaging by incubating the pads in CSM with the final desired drug concentration for 2-4 hours. The volume of the pad was included in calculating the final drug concentration. Pads for control cells were incubated in the same manner with the appropriate volume of DMSO. Pads were removed from the CSM-drug solution, briefly dried on a kimwipe, and then placed on cells. Prior to drug treatment, cells were grown in YPD without drug and were only exposed to the drug in the pad. To monitor growth, cells expressing the nuclear marker H2B-mCherry were imaged immediately after adding the drugged pads on a widefield Nikon ECLIPSE Ti2 inverted microscope with a Plan Apo 20x air objective (NA 0.75; Nikon Instruments), a sCMOS pco.edge camera (Excelitas Technologies), and a X-Cite XYLIS LED Illumination System (Excelitas Technologies) controlled by NIS-Elements software (Nikon Instruments). A single image was acquired every 5 minutes for 10 hours using an exposure time of 20 ms at 5% power (excitation 561 nm) to excite H2B-mCherry. To measure the effect of low does LatB (625 nM) on bud growth, single confocal stacks and time series were acquired of drug treated cells expressing the cytosolic marker 3xmCherry using the same spinning disk confocal setup as described above.

### Imaging phalloidin-stained F-actin

*act1^V75I^* cells were grown in YPD as for live imaging experiments. Cells were fixed by adding formaldehyde (CAS # 50-00-0, ThermoFisher) at a final concentration of 3.7% to 500 µl cells in YPD. To test the effect of LatB treatment on F-actin, cells were first treated with 50 µM or 625 nM LatB, or the carrier control (DMSO), for three hours prior to fixation. Cells were fixed for 40 minutes at 24°C with agitation. Following fixation, cells were rinsed three times with PBS and resuspended in 30 µl PBS with 0.1% Trition X-100. The Alexa Flour^TM^ 488 phalloidin (A12379, ThermoFisher) stock was prepared in anhydrous DMSO (66 µM final concentration) and 1.5 µl was added to 30 µl cells. Cells were incubated with phalloidin at 24°C with agitation for 2 hours in the dark, rinsed once with 100 µl PBS, mounted in SlowFade™ Glass Soft-Set Antifade Mountant (S36917, ThermoFisher), and imaged immediately. We found that more rinses with PBS or allowing the samples to sit at room temperature for more than one hour prior to imaging decreased the image quality. Fixed and stained cells were imaged on the same spinning disk confocal system used for live cell imaging described above. The entire cell volume was acquired using 75 Z-slices (at 0.2 μm step intervals). Exposure times of 50 ms at 50% laser power (excitation 488 nm) were used.

### Image analysis

The cytosolic marker 3xmCherry was used to segment and measure cell volumes of wild-type cells, cells treated with 625 nM LatB, cells expressing Pak1-GFP, or cells expressing Pak1ΔC-GFP. 3D segmentation and volume measurements were done in NIS-Elements General Analysis 3 software (GA3, Nikon Instruments).

Kymographs used to monitor bud growth and mNG-Tpm1 accumulation at the bud neck, to track GFP-Sec4 tagged vesicles, or to track For1-3xmNG and Pak1-mCherry, were generated in FIJI (Schindelin et al., 2012).

To measure total cable lengths in phalloidin stained cells, confocal Z-stacks were first denoised using DenoiseAI and background subtracted in NIS Elements (Nikon Instruments). Actin cables in processed Z-stacks were traced in 3D from the bud neck to the end of the cable using the Simple Neurite Tracer (SNT) plugin in FIJI (Arshadi et al., 2021). The total cable length was calculated by summing the lengths of all cables associated with a given bud neck.

The mNG-Tpm1 signal associated with cables in the bud was measured from confocal time series of cells expressing mNG-Tpm1 and the cytoplasmic marker, 3xmCherry. All image processing and measurements were carried out in NIS-Elements General Analysis 3 (GA3, Nikon Instruments). To remove the mNG-Tpm1 cytosolic background signal, confocal Z-stacks were background subtracted to set signal outside of cells to zero, cropped to include individual cells, and the signal intensity in the 488 nm (mNG-Tpm1) and 561 nm (3xmCherry) channels were measured for the individual cells. Then, the fold difference in signal intensity was calculated and the dimmer channel (488 nm) was multiplied by this difference such that the mean signal per cell in both channels, 488 nm and 561 nm, was the same. Finally, the signal in the 561 nm channel was subtracted from the 488 nm channel for each pixel in the Z-stack. The mean signal intensity in the processed 488 nm channel was measured from each 3D-segmented bud. Reported is the CV (standard deviation/mean) of this signal across all sibling buds, either for a single timepoint or after averaging across 10 time points (10 minutes).

The Pak1-GFP or Pak1ΔC-GFP signal was measured in bud volumes segmented using the 3xmCherry cytosolic marker in NIS-Elements General Analysis 3 software (GA3, Nikon Instruments). Reported are mean GFP intensities in each bud relative to bud volumes.

### Theoretical model of intracellular vesicle transport

We develop a mean-field biophysical model of intracellular vesicle transport to identify the key factors regulating vesicle delivery to multiple buds in *A. pullulans*. The framework approximates the shape of *A. pullulans* as an ellipsoid, characterized by major axis ℓ_x_, and minor axis ℓ_y_. Vesicles are synthesized in the cytoplasm at a rate k^&^ and diffuse through the cytoplasm with a diffusion coefficient D^v^. Vesicles are recruited to each bud through actin cables that emanate from the bud necks. Vesicles attach to cable networks at a rate given by the vesicle recruitment rate parameter k_on_. At the continuum level, the local vesicle recruitment rate by actin cables is proportional to the normalized actin cable intensity, whose spatial distribution is assumed to follow an exponential decay with distance from the bud neck, with length scale given by β. The spatiotemporal dynamics of the cytoplasmic vesicle concentration C_v_ is therefore described by the following mass conservation equation:

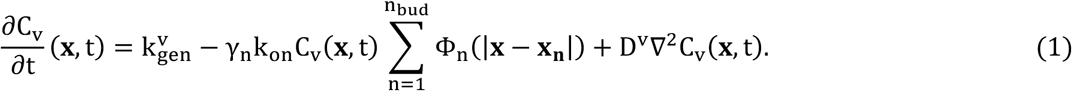

The term on the left-hand-side (LHS) of Eq. (1) represents the rate-of-change of cytoplasmic vesicle density. The three terms on the right-hand-side (RHS) correspond to homogeneous vesicle generation, vesicle recruitment by actin cables, and vesicle diffusion, respectively. The function Φ_n_ represents the normalized actin cable intensity linked to bud n, while x_n_ denotes its position vector. To isolate the systematic effects of bud placement asymmetry on vesicle recruitment rates and eliminate stochastic variability, we introduce the parameter constants γ_n_, ensuring equal actin cable intensity across all buds. These parameters satisfy the condition:

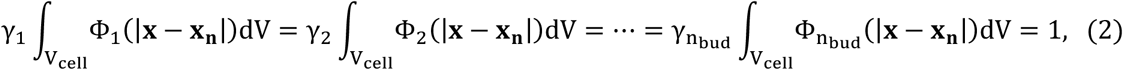

where V_cell_ represents the cell volume. The number of vesicles recruited by bud n is determined by the total vesicles captured across the cytoplasm by actin cables emanating from its neck:

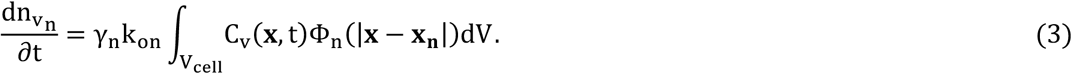

We impose a no-flux boundary condition for vesicle diffusion at the cell membrane:

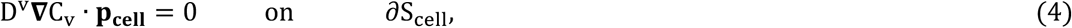

where **p_cell_** denotes the outward unit normal vector to the cell surface, and ∂**S_cell_** represents the cell boundary. We solve Eqs. (1–3) using the finite volume method, approximating spatial derivatives with second-order central differences, and computing time integrals using the explicit Euler method with a time step of 10^-4^. The computational domain, of size L_x_ = 10.5 μm, L_y_ = 6.6 μm, L_z_ = 6.6 μm, is discretized with equally spaced collocation points, with N_x_ = 200, N_y_ = 120, and Nz = 120 collocation points in the x-, y– and z-directions, respectively. Here, the x-axis is aligned with the major axis of the cell.

### Model parameters

The ranges of different model parameters have been estimated from images (cell dimensions, actin length scale, actin cable number and hence recruitment rate) or expectations from the cell biology literature (vesicle diffusion constants).

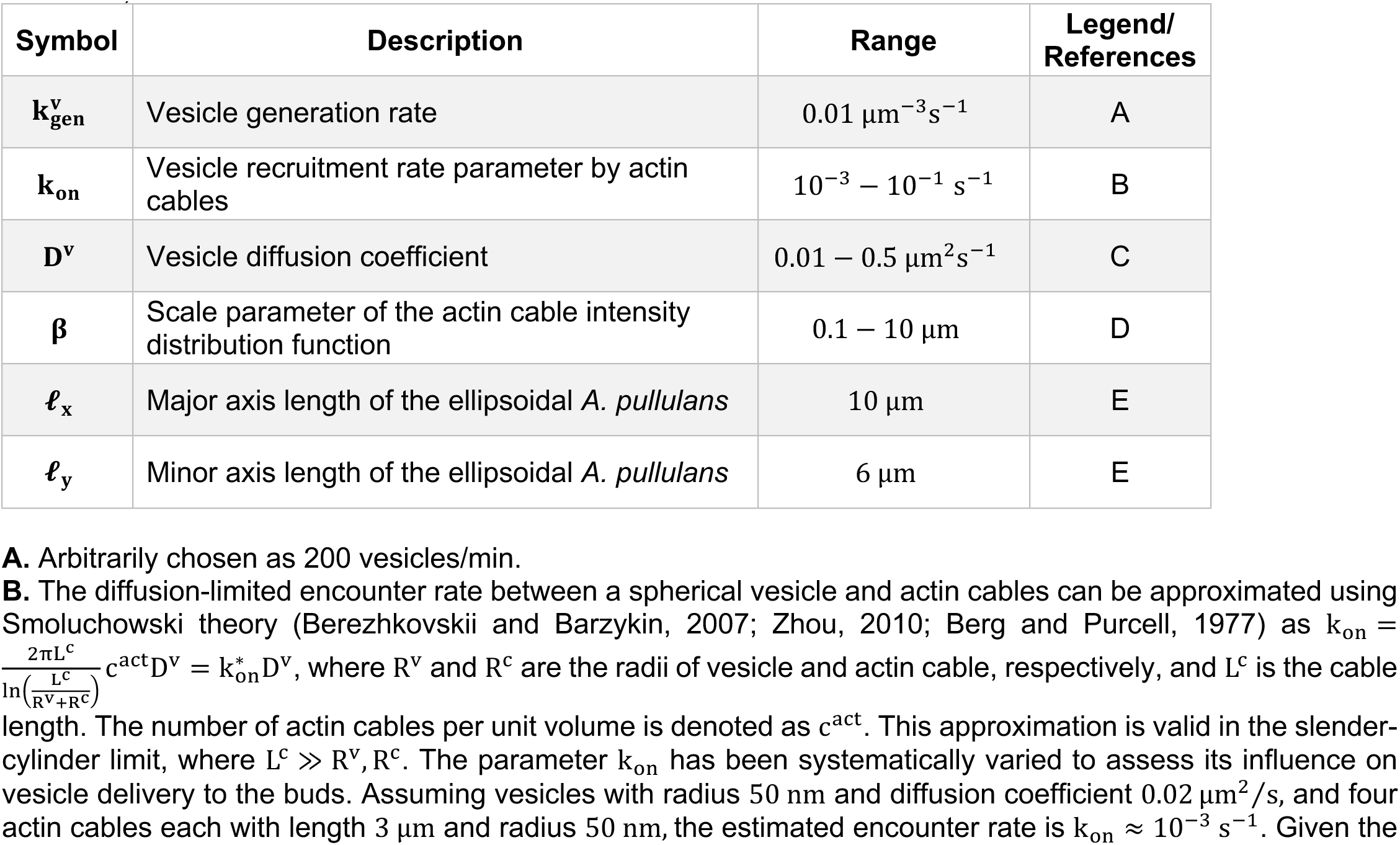
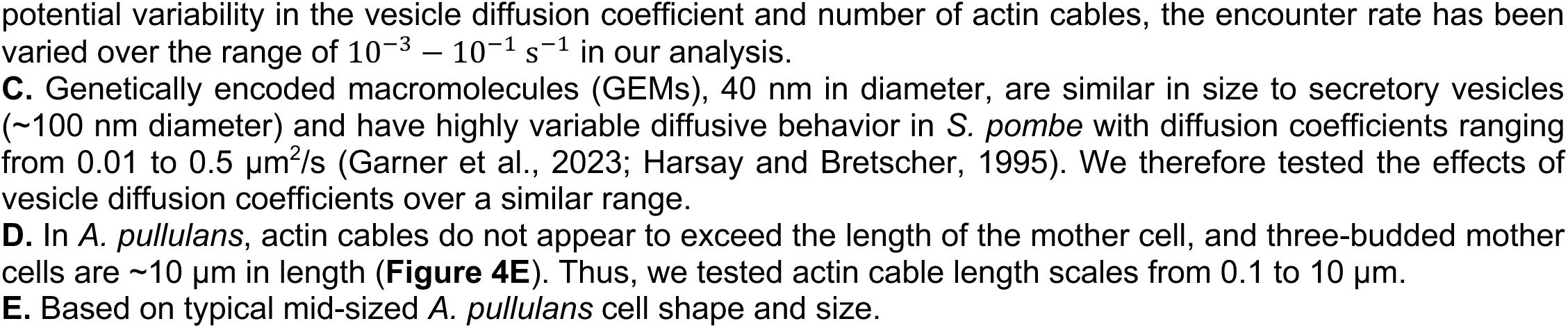

| Symbol | Description | Range | Legend/References |
| --- | --- | --- | --- |
| $k_{\text{gen}}^v$ | Vesicle generation rate | $0.01 \mu\text{m}^{-3} \text{s}^{-1}$ | A |
| $k_{\text{on}}$ | Vesicle recruitment rate parameter by actin cables | $10^{-3} - 10^{-1} \text{s}^{-1}$ | B |
| $D^v$ | Vesicle diffusion coefficient | $0.01 - 0.5 \mu\text{m}^2 \text{s}^{-1}$ | C |
| $\beta$ | Scale parameter of the actin cable intensity distribution function | $0.1 - 10 \mu\text{m}$ | D |
| $\ell_x$ | Major axis length of the ellipsoidal *A. pullulans* | $10 \mu\text{m}$ | E |
| $\ell_y$ | Minor axis length of the ellipsoidal *A. pullulans* | $6 \mu\text{m}$ | E |

**A.** Arbitrarily chosen as 200 vesicles/min.
**B.** The diffusion-limited encounter rate between a spherical vesicle and actin cables can be approximated using Smoluchowski theory (Berezhkovskii and Barzykin, 2007; Zhou, 2010; Berg and Purcell, 1977) as $k_{\text{on}} = \frac{2\pi L^c}{\ln(\frac{L^c}{R^v + R^c})} c^{\text{act}} D^v = k_{\text{on}}^* D^v$ , where $R^v$ and $R^c$ are the radii of vesicle and actin cable, respectively, and $L^c$ is the cable
length. The number of actin cables per unit volume is denoted as $c^{\text{act}}$ . This approximation is valid in the slender-cylinder limit, where $L^c \gg R^v, R^c$ . The parameter $k_{\text{on}}$ has been systematically varied to assess its influence on vesicle delivery to the buds. Assuming vesicles with radius 50 nm and diffusion coefficient $0.02 \mu\text{m}^2/\text{s}$ , and four actin cables each with length 3 $\mu\text{m}$ and radius 50 nm, the estimated encounter rate is $k_{\text{on}} \approx 10^{-3} \text{s}^{-1}$ . Given the
potential variability in the vesicle diffusion coefficient and number of actin cables, the encounter rate has been varied over the range of $10^{-3} - 10^{-1} \text{ s}^{-1}$ in our analysis.
**C.** Genetically encoded macromolecules (GEMs), 40 nm in diameter, are similar in size to secretory vesicles (~100 nm diameter) and have highly variable diffusive behavior in *S. pombe* with diffusion coefficients ranging from 0.01 to $0.5 \mu\text{m}^2/\text{s}$ (Garner et al., 2023; Harsay and Bretscher, 1995). We therefore tested the effects of vesicle diffusion coefficients over a similar range.
**E.** Based on typical mid-sized *A. pullulans* cell shape and size.

### Growth assay

To compare cell growth of *A. pullulans* strains, a single colony was inoculated into 5 ml of YPD (2% glucose) and grown at 24°C for 48 h. Cultures were serially diluted five times in sterile deionized water in a 96-well plate and transferred onto solid YPD plates using a pin-frogger. Plates were grown for 48 h at 24°C and imaged on an Amersham Imager 680 (General Electric Company) using the colorimetric Epi-white settings.

### Statistical analysis

All statistical analysis was done using GraphPad Prism. Unless indicated otherwise, data distributions were assumed to be normal, but this was not formally tested. Statistical comparison between indicated conditions was conducted using the two-sided Student’s t test or one-way ANOVA, as indicated in the figure legends. After running an ANOVA, the Tukey test was used to compare the mean with every other mean. Differences were considered significant if the p-value was <0.05.

## SUPPLEMENTAL MATERIAL

Supplemental Figure 1 shows growth assays for each of the strains built for this study expressing fluorescently tagged or mutated proteins. Supplemental Figure 2 shows examples of phalloidin stained cells treated with different concentrations of LatB. Supplemental Table 1 provides a list of all strains used in this study. Supplemental Table 2 provides a list of plasmids generated in this study. Supplemental Table 3 provides a list of all primers used to amplify DNA to transform *A. pullulans*.

## DATA AVAILABILITY

The data generated in this study are available from the corresponding author upon reasonable request.

## ABBREVIATIONS

DNA: Deoxyribonucleic acid
doURA: media lacking uracil
ER: Endoplasmic reticulum
ERES: ER exit sites
G418: Geneticin
GEF: Guanine nucleotide exchange factor
GFP: Green Fluorescent Protein
Hyg: hygromycin B
mNG: mNeonGreen
Nat: nourseothricin

## ACKNOWLEDGEMENTS

We would like to thank Avril Wang, Claudia Petrucco, and Yiqiao Zheng for their helpful comments on this work. This work was funded by NIH/NIGMS grant R35GM122488 to DJL, and NIH/NCI grants P01CA254849 and U54CA268069 to D.J.O.

## AUTHOR CONTRIBUTIONS

Conceptualization, review, and editing manuscript— A.C.E. Wirshing, R. Alonso-Matilla, D. J. Odde, and D.J. Lew. Data curation, investigation, methodology, visualization, validation, and formal analysis— A.C.E. Wirshing, R. Alonso-Matilla, M. Yan, S. Khalid, and A. Colarusso. Drafting of manuscript— A.C.E. Wirshing. Project administration, supervision, and resources— D. J. Odde and D.J. Lew.

## SUPPLEMENTAL FIGURE CAPTIONS

**Supplemental Figure 1:**
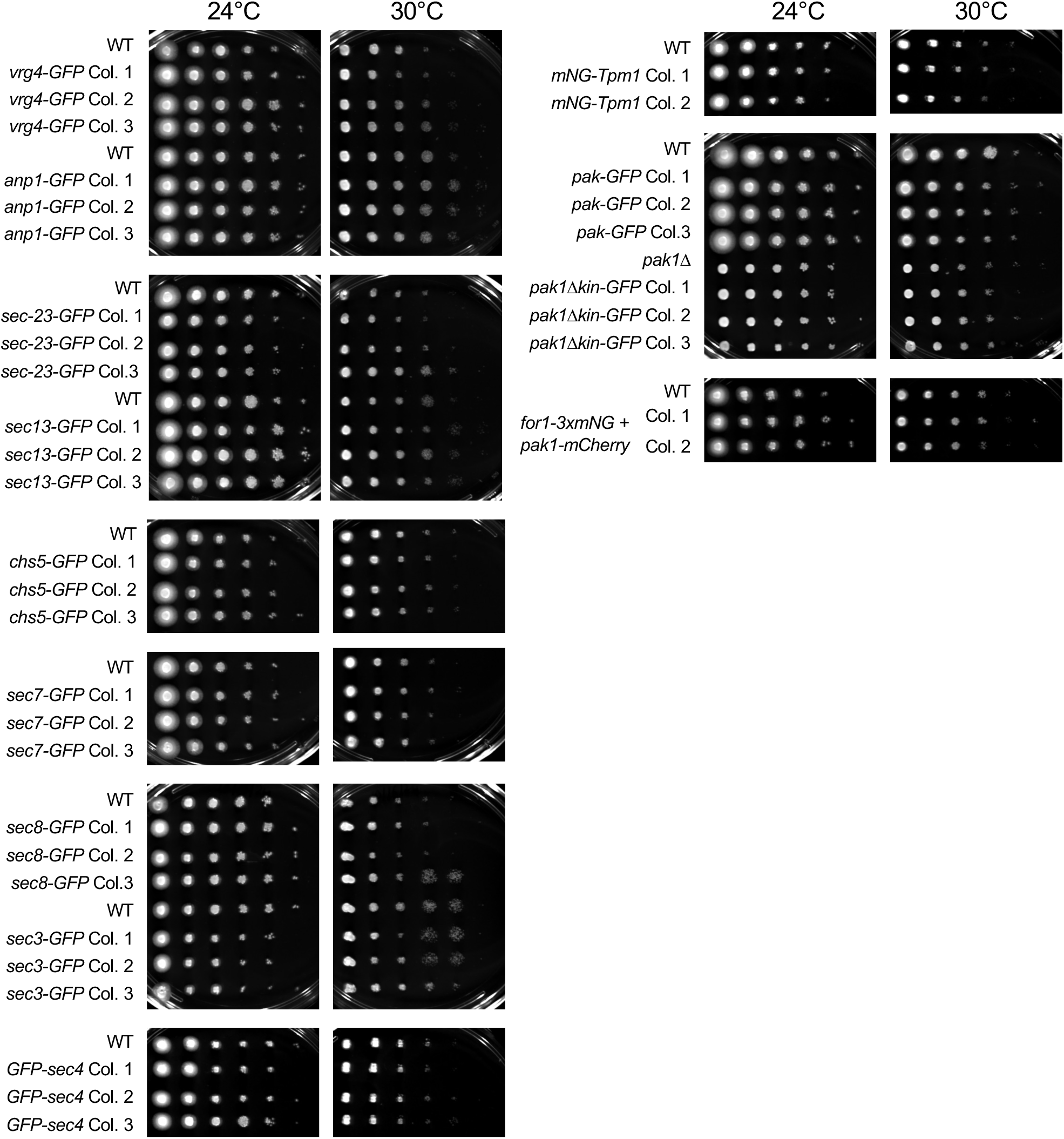
Growth assays of 10-fold serial dilutions of the indicated strains plated on YPD and grown for 48 hours at the indicated temperature. Three independent transformants (Colony 1-3) are shown.

**Supplemental Figure 2.**
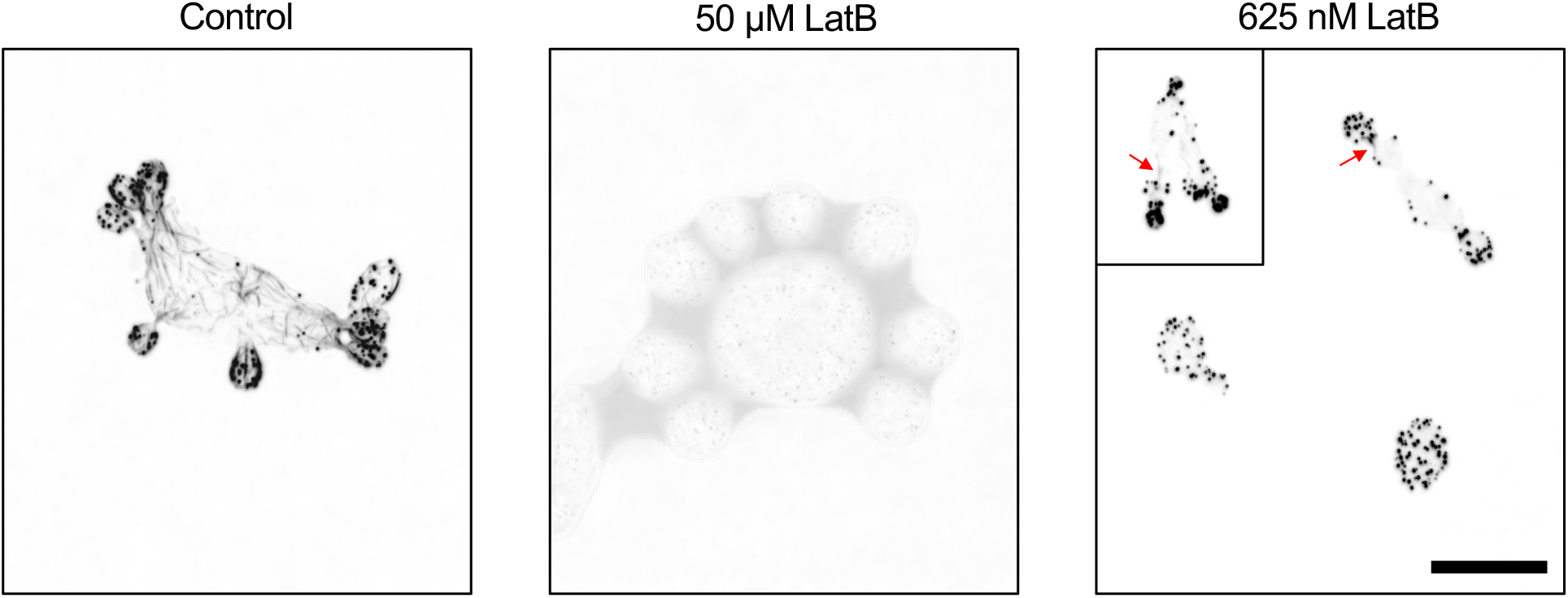
: Effect of LatB on F-actin. Confocal maximum projection images of *act1*^V75I^ cells treated with 50 µM LatB, 625 nM LatB, or vehicle control (DMSO) and then fixed and stained with Alexa488-phalloidin. Images are inverted to highlight dim actin cables. With 50 µM LatB, no clear F-actin structures are seen. With a low dose of LatB, 625 nM, actin patches are depolarized and faint actin cables are only occasionally visible (red arrow). Scale bar, 5 µm.

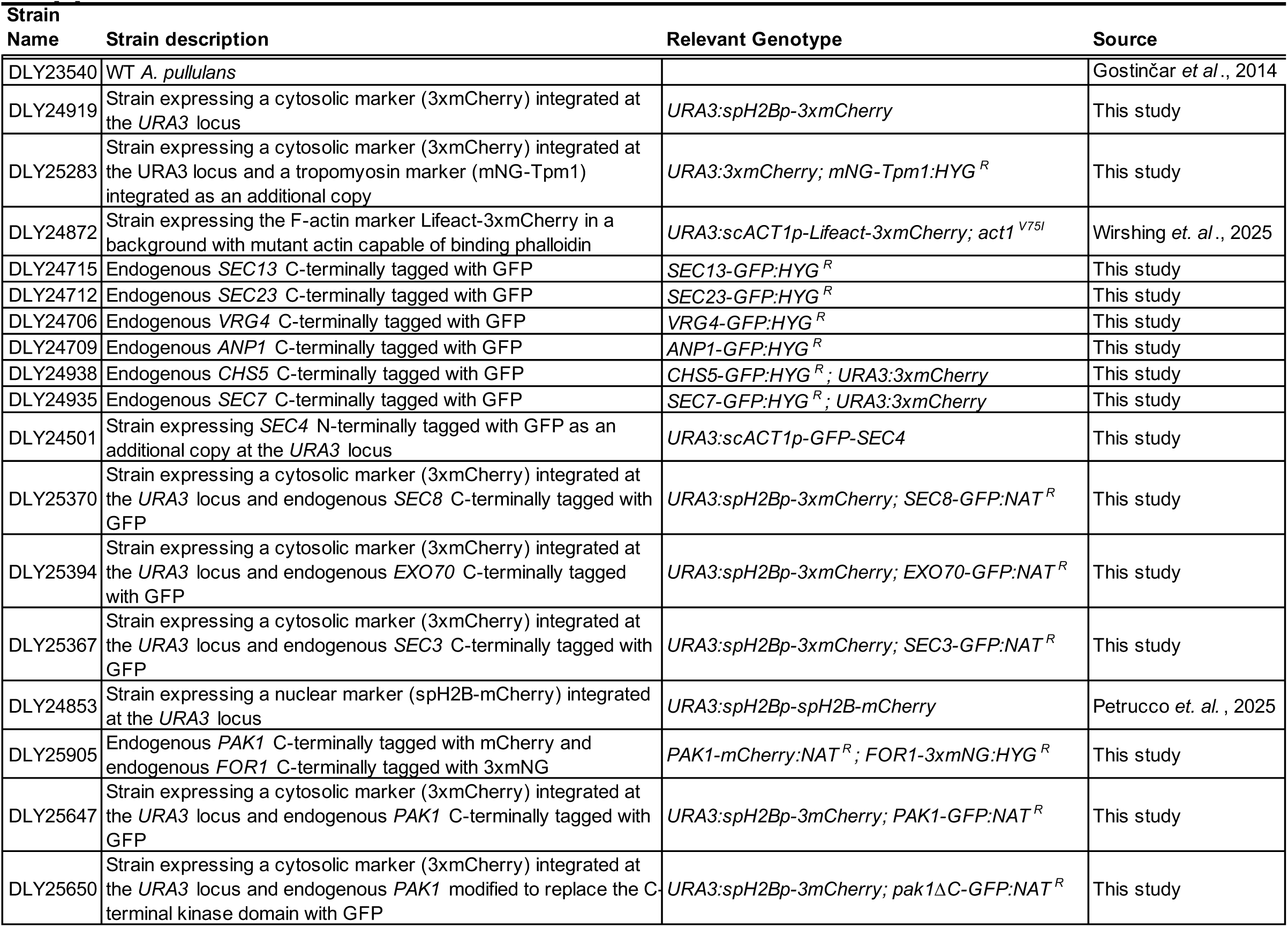
Supplemental Table 1.

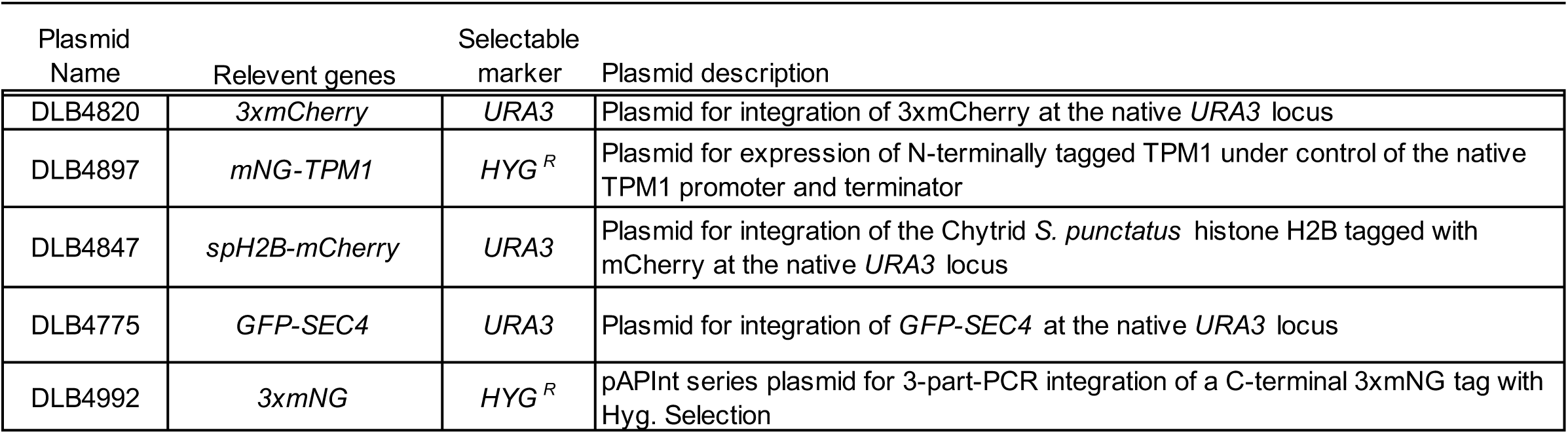
Supplemental Table 2.

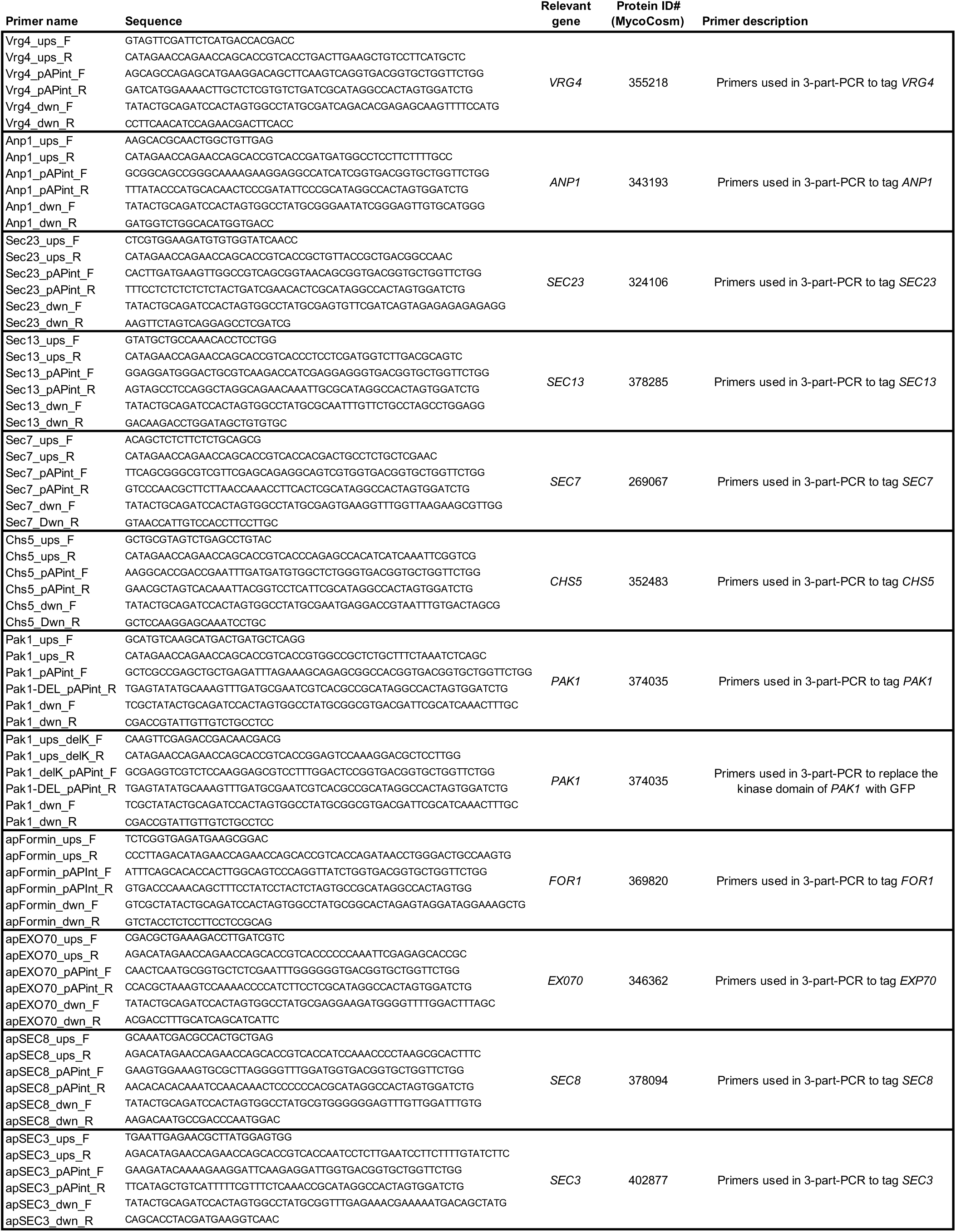
Supplemental Table 3.

## Notes

### Competing Interest Statement

The authors have declared no competing interest.

